# Zic1 advances epaxial myotome morphogenesis to cover the neural tube via Wnt11r

**DOI:** 10.1101/2021.07.12.452069

**Authors:** Ann Kathrin Heilig, Ryohei Nakamura, Atsuko Shimada, Yuka Hashimoto, Yuta Nakamura, Joachim Wittbrodt, Hiroyuki Takeda, Toru Kawanishi

## Abstract

The dorsal axial muscles, or epaxial muscles, are a fundamental structure covering the spinal cord and vertebrae, as well as mobilizing the vertebrate trunk. To date, mechanisms underlying the morphogenetic process shaping the epaxial myotome are largely unknown. To address this, we used the medaka *zic1/zic4*-enhancer mutant *Double anal fin* (*Da*), which exhibits ventralized dorsal trunk structures resulting in impaired epaxial myotome morphology and incomplete coverage over the neural tube. In wild type, dorsal dermomyotome (DM) cells, progenitors of myotomal cells, reduce their proliferative activity after somitogenesis and subsequently form unique large protrusions extending dorsally, potentially guiding the epaxial myotome dorsally. In *Da*, by contrast, DM cells maintain the high proliferative activity and form mainly small protrusions. By combining RNA- and ChIP-sequencing analyses, we revealed direct targets of Zic1 which are specifically expressed in dorsal somites and involved in various aspects of development, such as cell migration, extracellular matrix organization and cell-cell communication. Among these, we identified *wnt11r* as a crucial factor regulating both cell proliferation and protrusive activity of DM cells. We propose that the dorsal movement of the epaxial myotome is guided by DM cells and that Zic1 empowers this activity via Wnt11r to achieve the neural tube coverage.

## Introduction

Active locomotion, which is powered by skeletal muscles in vertebrates, is critical for animals to survive. Vertebrate skeletal muscles consist of axial muscles (head, trunk and tail muscles) and appendicular muscles (limb muscles). Axial muscles first arose in the chordates to stabilize and enable side-to-side movement of the body axis. In jawed vertebrates, the subdivision of axial muscles into epaxial (dorsal) and hypaxial (ventral) muscles led to an increased range of movement: dorsoventral undulation in fish and lateral movements in terrestrial vertebrates (Glass and Goodrich, 1960; Romer and Parsons, 1986; Fetcho, 1987; Clifton, 2002; Sefton and Kardon, 2019). Among these muscles, epaxial muscles are characterized by their unique anatomical structure which extends dorsally and surrounds the vertebrae. This morphology also ensures mechanical support and protection of the vertebrae and the spinal cord inside. While we have a detailed understanding of how the myotome, precursors of epaxial and hypaxial muscles, differentiates from the somites (Kalcheim and C, 1999; Gros, Scaal and Marcelle, 2004; Hollway and Currie, 2005), we only begin to understand the cellular and molecular mechanisms of the subsequent morphogenetic processes generating the epaxial muscles. Previous studies in rats and mice suggested that myocytes of the epaxial myotome do not actively migrate dorsally but are guided by external forces (Deries, Schweitzer and Duxson, 2010; Deries *et al*., 2012). However, what exerts such forces to drive extension of epaxial myotome is still unclear.

Fish have been extensively utilized to study myotome development thanks to the transparency of their bodies throughout embryonic development (Nguyen *et al*., 2017; Ganassi *et al*., 2018). Additionally, their epaxial trunk muscles have a simple structure consisting of only one anatomical unit (reviewed in (Sefton and Kardon, 2019)). Like other vertebrates, fish myotomes, on either side of the neural tube, extend dorsally after somite differentiation and eventually cover the neural tube by the end of embryonic development (Figure 1A). The spontaneous medaka (*Oryzias latipes*) mutant *Double anal fin* (*Da*) displays a particular epaxial myotome morphology, in which the dorsal ends of the left and right epaxial myotome fail to extend sufficiently and thus do not cover the neural tube at the end of embryonic development. Previous studies demonstrated that the dorsal trunk region of the *Da* mutant is transformed into the ventral one, including not only the myotome but also the body shape, skeletal elements, pigmentation and fin morphology (Figure 1B, D) (Ishikawa, 1990; Ohtsuka *et al*., 2004). Given the unique morphological features, the medaka *Da* mutant is an excellent model to study the morphogenesis of epaxial myotome. Genetic analysis of the *Da* mutant revealed that this phenotype is due to a dramatic reduction of the expression of the transcription factors *zic1* and *zic4* in the dorsal somites (Figure 1C, E), and identified *zic*1/*zic4* as master regulators of trunk dorsalization (Ohtsuka *et al*., 2004; Kawanishi *et al*., 2013). The down-regulation of *zic1/zic4* specifically in the dorsal somites is caused by the insertion of a large transposon, disrupting the dorsal somite enhancer of *zic1/zic4* (Moriyama *et al*., 2012; Inoue *et al*., 2017). While the function and downstream targets of Zic1 and Zic4 have been studied in the nervous system (Aruga and Millen, 2018), the molecular mechanism of how these Zic genes control dorsal trunk morphogenesis has not been investigated so far.

**Figure 1:**
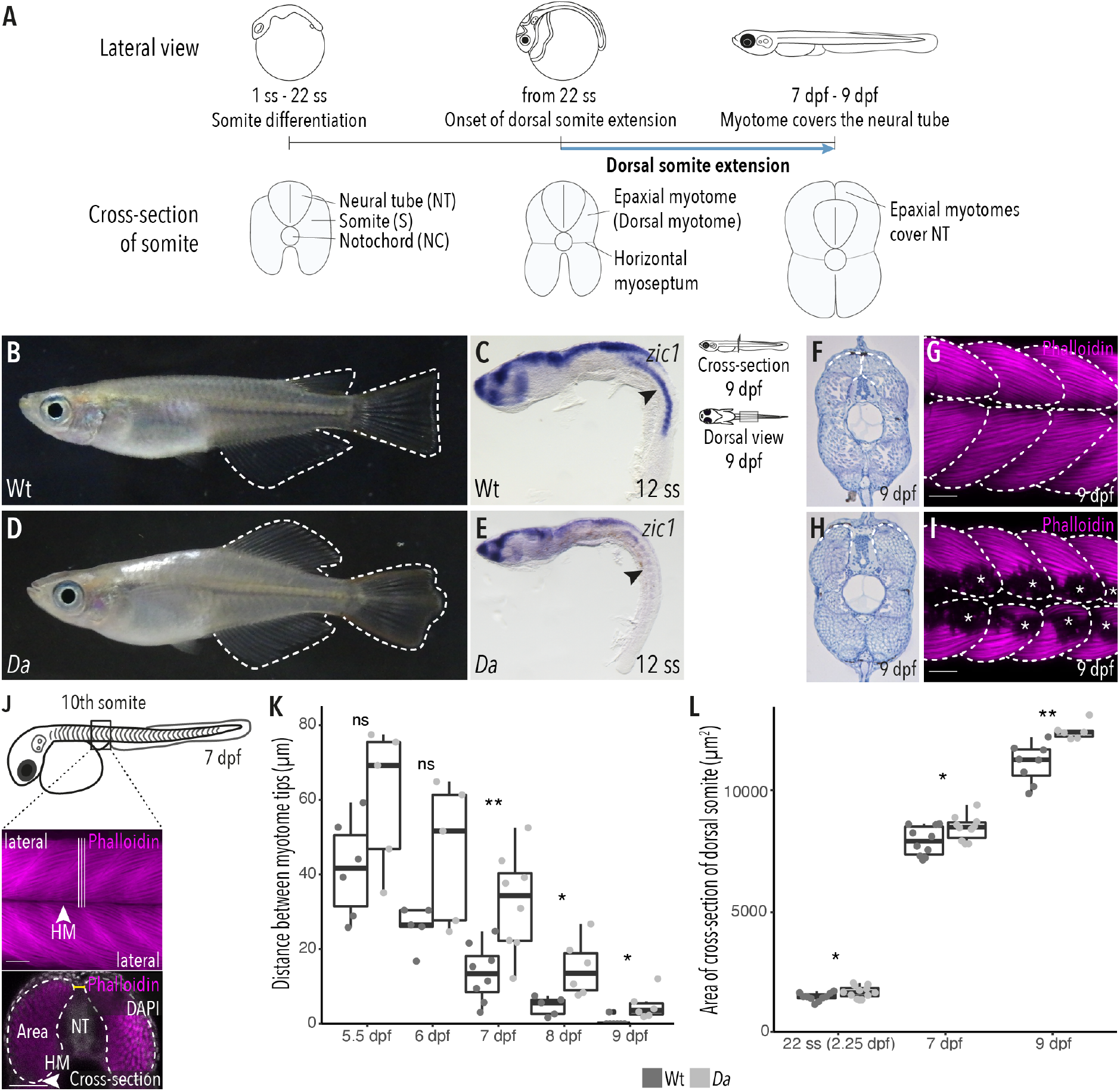
The epaxial myotome of the *Da* mutant fails to cover the neural tube at the end of embryonic development. (A) Schematic representation of dorsal somite extension which results in the full coverage of the neural tube by the epaxial myotomes at the end of embryonic development. (B) Lateral view of adult Wt medaka. Dorsal, caudal and anal fins are outlined. (C) Lateral view of whole-mount *in situ* hybridization against *zic1* in a 12 ss (1.7 dpf, stage 23) Wt embryo. *Zic1* expression can be observed in the brain, neural tissues and the dorsal somites (arrowhead). (D) Lateral view of adult *Da* mutant. Dorsal, anal and caudal fins are outlined. The dorsal trunk region resembles the ventral trunk region. (E) Lateral view of whole-mount *in situ* hybridization against *zic1* of a 12 ss *Da* embryo. *Zic1* expression can be observed in the brain and the neural tissues, but is drastically decreased in the dorsal somites (arrowhead). (F, H) Cross-sections of tail regions of hematoxylin stained 9 dpf embryos. Dorsal ends of myotomes are outlined. (F) In Wt, the left and the right myotome come in close contact at the top of the neural tube and form a gapless muscle layer. (H) In the *Da* mutant, the left and right myotome fail to come in contact at the top of the neural tube. (G, I) Dorsal view of whole-mount Phalloidin (magenta) immunostaining labeling the myotome of (G) Wt and (I) *Da* embryos. Epaxial myotome is outlined, asterisks label melanophores. (J) Schematic representation of measurements to analyze the distance between the left and the right dorsal tip of the myotome (yellow) and the cross-sectional area of the dorsal myotome. For each measurement, three consecutive optical cross sections of the 10^th^ somite were analyzed and averaged. (K) Analysis of the distance between the left and right tip of the dorsal myotome 5.5 dpf – 9 dpf (n = 6 Wt embryos 5.5 dpf (stage 35), n = 5 *Da* embryos 5.5 dpf, n = 5 Wt embryos 6 dpf (stage 36), n = 5 *Da* embryos 6 dpf, n = 8 Wt embryos 7 dpf, n = 8 *Da* embryos 7 dpf, n = 5 Wt embryos 8 dpf (stage 38), n = 6 *Da* embryos 8 dpf, n = 7 Wt embryos 9 dpf, n = 6 *Da* embryos 9 dpf, median, first and third quartiles, ^ns^P_5.5 dpf_ = 0.097, ^ns^P_6 dpf_ = 0.075, **P_7 dpf_ = 0.0047, *P_8 dpf_ = 0.019 *P_9 dpf_ = 0.034). (L) Analysis of cross-sectional area of the dorsal somites at 22 ss (2.25 dpf, stage 26; n = 10 somites of 5 Wt embryos, n = 12 somites of 6 *Da* embryos), 7 dpf (n = 10 somites of 5 Wt embryos, n = 10 somites of 5 *Da* embryos) and 9 dpf (n = 8 somites of 4 Wt embryos, n = 6 somites of 3 *Da* embryos) (median, first and third quartiles, *P_22 ss_ = 0.038, *P_7 dpf_ = 0.044, **P_9 dpf_ = 0.0019). Anterior = left, HM = horizontal myoseptum, NT = neural tube, scale bar = 50 μm.

Here we describe the morphogenetic process of the formation of the epaxial myotome of the back, which we termed “dorsal somite extension” (Figure 1A). By *in vivo* time-lapse imaging we uncovered its cellular dynamics; during dorsal somite extension, dorsal dermomyotome (DM) cells reduce their proliferative activity and subsequently form unique large protrusions extending dorsally, potentially guiding the epaxial myotome dorsally. In the *Da* mutant, by contrast, DM cells keep their high proliferative activity and mainly form small protrusions. Mechanistically, we identify a Zic1 downstream-target gene, *wnt11r*, as a crucial factor for dorsal somite extension. We demonstrate that Wnt11r regulates cellular behavior of dorsal DM cells by promoting protrusion formation and negatively regulating proliferation.

## Results

Our previous study showed that *zic1* and *zic4* expression starts at embryonic stages and persists throughout life (Kawanishi *et al*., 2013). Phenotypic analysis of homozygous adult *Da* mutants implies long-term participation of Zic-downstream genes in the formation of dorsal musculatures, which eventually affects the external appearance of the fish adult trunk. Here, we examined the initial phase of this long-term dorsalization process. In the following of the study, we will focus on *zic1*, since *zic1* and *zic4* are expressed in an identical fashion with overlapping functions in trunk dorsalization of medaka, and *zic4* is expressed slightly weaker than *zic1* (Moriyama *et al*., 2012; Kawanishi *et al*., 2013).

### The dorsal myotome of the *Da* mutant fails to cover the neural tube

In wild type (Wt) medaka, the dorsal ends of the myotomes first came in contact at 7 days post fertilization (dpf, stage 37) and formed the tight, thick myotome layer covering the neural tube at the end of embryonic development (9 dpf, stage 39) (Figure 1F-G, Figure 1 - figure supplement 1A-F). In the ventralized *Da* mutant, however, the dorsal ends of the myotomes did not extend sufficiently and failed to cover the neural tube at the end of embryonic development (Figure 1H-I, Figure 1 - figure supplement 1G-L). The ends of the ventralized dorsal myotome in the mutant displayed a round shape (not pointed as found in Wt) which resembled the morphology of the ventral myotome.

We wondered if there are other morphological differences between Wt dorsal myotome and the ventralized dorsal myotome of the *Da* mutant. Indeed, the cross-sectional area in the *Da* mutant was significantly larger compared to Wt (Figure 1L). Possible explanations for a larger cross-sectional area in the *Da* mutant could be a larger myofiber diameter or a higher number of myofibers which make up the myotome. We measured the diameter of dorsal myotome muscle fibers in Wt and *Da* mutant embryos but could not observe a difference, suggesting that dorsal myotome of the *Da* mutant has a higher number of myotomal cells (Figure 1 - figure supplement 1M).

### Proliferative activity of the dorsal DM cells is enhanced in the *Da* mutant

In fish, as in other vertebrates, the DM gives rise to muscle precursor cells which ultimately differentiate into myofibers. In medaka the DM is a one cell-thick, Pax3/7-positive cell layer encompassing the myotome (Figure 2A-B’’) (Hollway *et al*., 2007; Abe *et al*., 2019). A high proliferative activity of the dorsal DM could explain a larger cross-sectional area of the dorsal myotome of the *Da* mutant. To test this, we performed immunohistochemistry against the mitotic marker phosphor-histone H3 (PH3) and the DM marker Pax3/7 on Wt and *Da* embryos (Figure 2C-E). In both Wt and *Da* embryos, PH3-positive cells were randomly distributed in the dorsal DM without obvious bias (Figure 2C-D). At the 12-somite stage (12 ss, 1.7 dpf stage 23), when *zic1* expression in the somites becomes restricted to the dorsal region (Kawanishi *et al*., 2013), the number of PH3-positive DM cells per dorsal somite was not significantly different between Wt and *Da*. Remarkably, from 16 ss (1.8 dpf, stage 24) onwards, the number of PH3-positive cells became reduced in the Wt, whereas in *Da*, no such reduction was observed (Figure 2E). At 35 ss (3.4 dpf, stage 30), PH3-positive cells increased both in the Wt and *Da*, but the mutant DM cells were more proliferative (Figure 2E). The number of PH3-positive cells in the ventral DM was not significantly different in Wt and *Da* embryos at 22 ss (2.75 dpf, stage 26) (Figure 2—figure supplement 1A). These results suggest that *zic1* reduces proliferative activity of the DM, which becomes evident following the confinement of its expression to the dorsal somite region.

**Figure 2:**
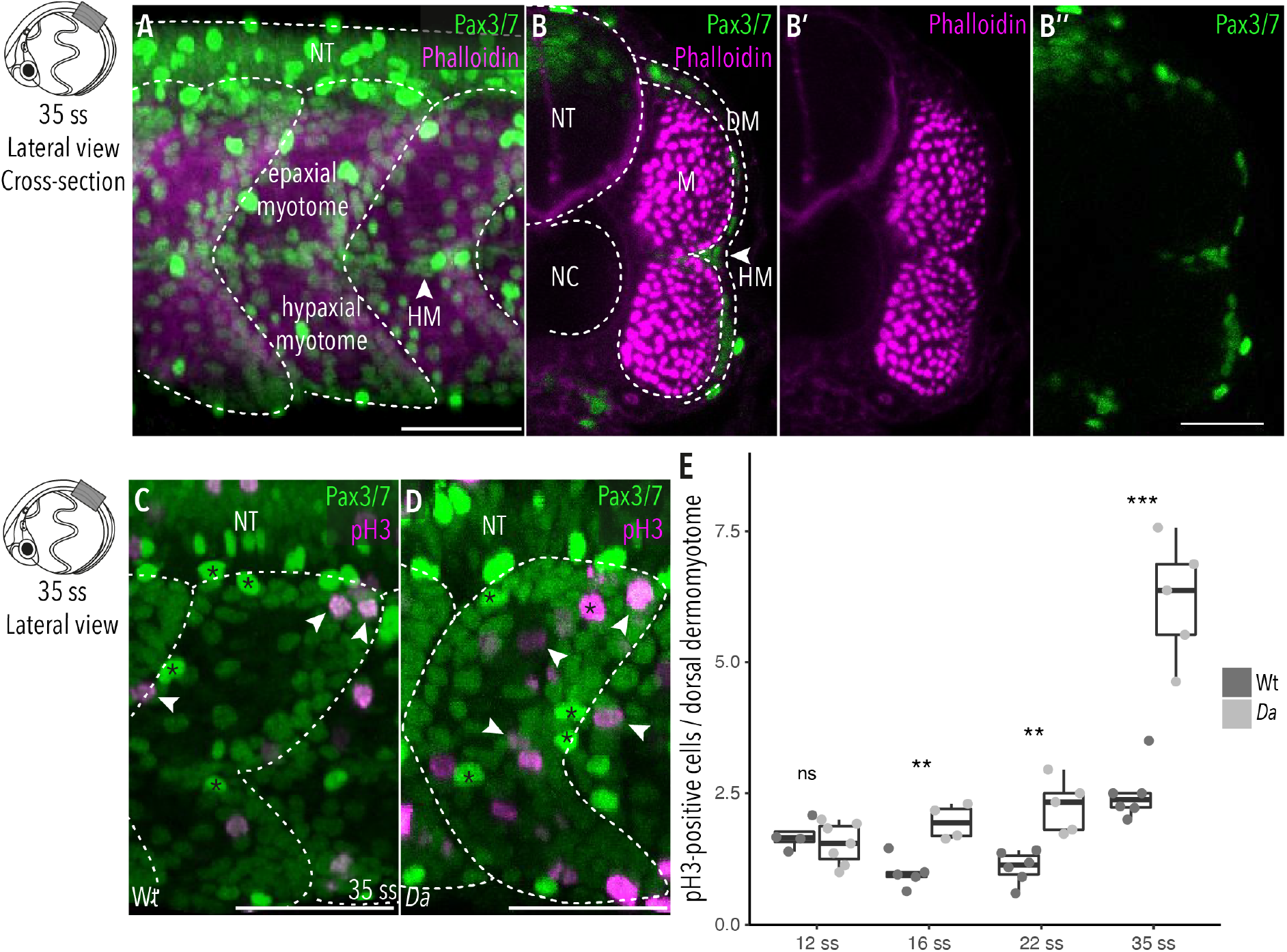
Wt dorsal DM cells show lower proliferative activity after the confinement of *zic1* expression to the dorsal somite. (A) Lateral view of 35 ss (3.4 dpf, stage 30) embryo, 10^th^ somite is positioned in the center. Pax3/7 (green) labels DM cells and Phalloidin (magenta) labels myotome. The horizontal myoseptum (HM) separates the myotome into epaxial myotome (dorsal) and hypaxial myotome (ventral). (B–B’’) Optical cross sections, myotome is labelled by Phalloidin (magenta) and encompassed by a one-cell thick layer of DM labelled with Pax3/7 (green). (C-D) Lateral view of (C) Wt and (D) *Da* 35 ss embryos. DM is labelled with Pax3/7 (green), mitotic active cells are PH3-positive (magenta, arrowheads (exemplary)). Asterisks exemplary mark neural crest cells which are highly Pax3/7-positive. (E) Quantification of PH3-positive cells in Wt and *Da* at 12 ss (n = 46.5 somites from 4 Wt embryos, n = 95 somites from 7 *Da* embryos), 16 ss (n = 54.5 somites from 5 Wt embryos, n = 42.5 somites from 4 *Da* embryos), 22 ss (n = 66 somites from 6 Wt embryos, n = 40.5 somites from 5 *Da* embryos,) and 35 ss (n = 49 somites from 6 Wt embryos, n = 47.5 somites from 5 *Da* embryos,) (median, first and third quartiles, ^ns^P_12 ss_ = 0.48, **P_16 ss_ = 0.0038, **P_22 ss_ = 0.0035, ***P_35 ss_ = 0.0008). Anterior = left, dorsal = up, DM = dermomyotome, HM = horizontal myoseptum, M = myotome, NT = neural tube, NC = notochord, scale bar = 50 μm.

### Wt dorsal DM cells form numerous large, motile protrusions at the onset of dorsal somite extension

The epaxial myotome, on either side of the neural tube, extends dorsally to cover the neural tube by the end of embryonic development. To examine the behavior of *zic1*-positive cells underlying this dorsal somite extension, we performed *in vivo* time-lapse imaging of dorsal somites using the transgenic line Tg(*zic1:GFP; zic4:DsRed*), which expresses GFP under the control of the *zic1* promoter and enhancers to visualize the dorsal somitic cells (Kawanishi *et al*., 2013) (hereafter called Tg(*zic1:GFP*) since the DsRed fluorescence was negligible in the following analyses). Intriguingly, around 22 ss and onwards, cells at the tip of the dorsal somites started to form numerous large protrusions extending dorsally towards the top of the neural tube (Figure 3A, Video 1). We defined the beginning of protrusion formation as the onset of dorsal somite extension. Close-up views of the time-lapse images (Figure 3A, Video 1) showed that these protrusions were motile and dynamically formed new branches at their dorsal tips (Figure 3J). Immunohistochemistry revealed that the protrusion-forming cells belong to the DM (Figure 3 - figure supplement1 A-A’’).

**Figure 3:**
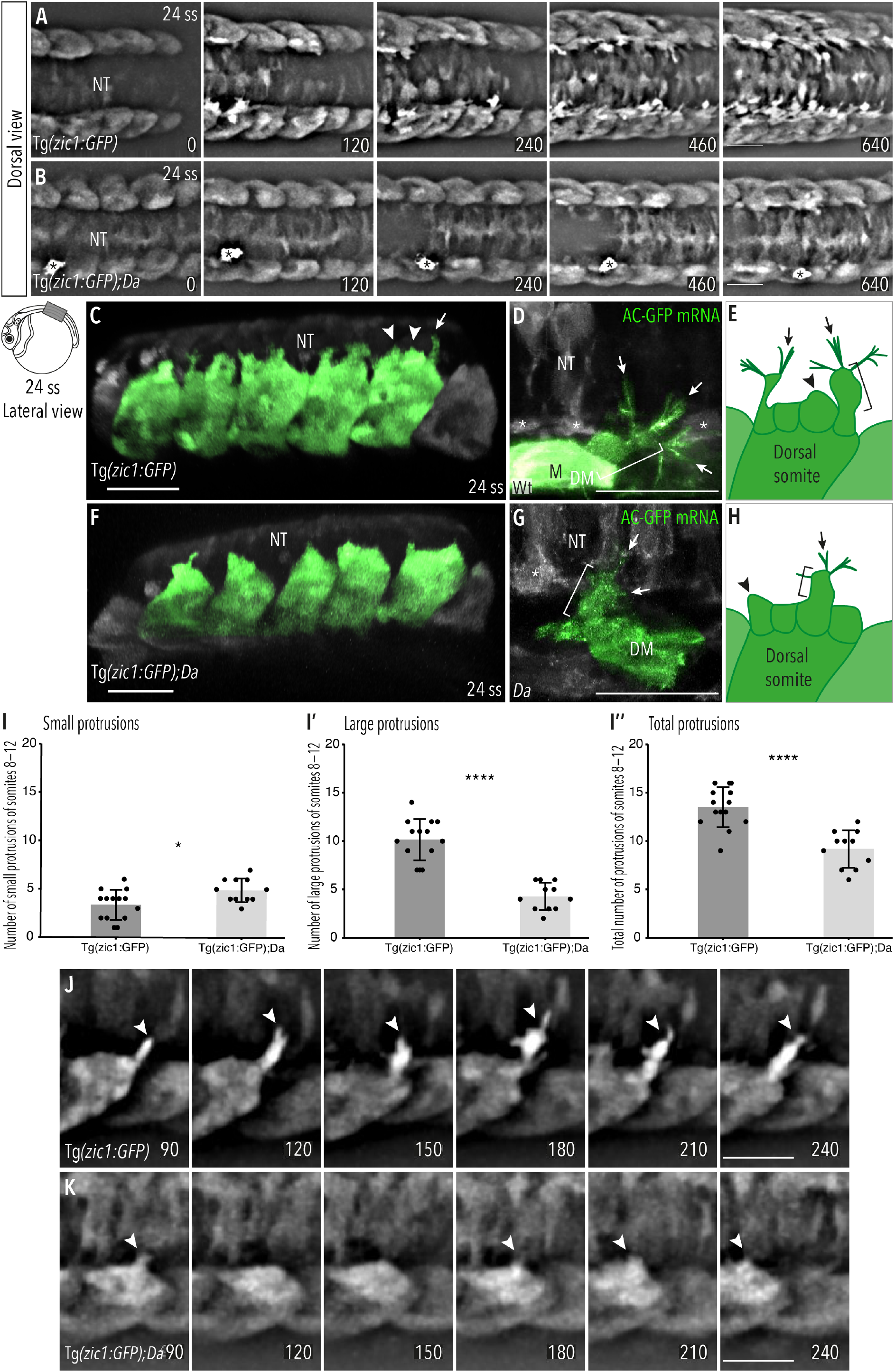
Wt dorsal DM cells form numerous large, motile protrusions at the onset of somite extension. (A-B) Dorsal view of time-lapse *in vivo* imaging of onset of dorsal somite extension at 24 ss (2.4 dpf, stage 27) of (A) Tg(*zic1:GFP*) and (B) Tg(*zic1:GFP);Da* embryos. 15^th^ somite is positioned in the center, z-stacks were imaged every 10 min, time is displayed in min. Asterisks indicate migrating melanophore. Scale bar = 50 μm. (C, F) Lateral view of 3D-reconstructions of *in vivo* imaging of (C) Tg(*zic1:GFP*) and (F)Tg(*zic1:GFP*);*Da*, dorsal somites are colored green. Arrowheads indicate small protrusions, arrow indicates large protrusion. Scale bar = 50 μm. (D, G) Dorsal view of 3D-reconstruction of *in vivo* imaging of large protrusion in (D) Wt and (G) *Da*. Embryos were injected with Actin-Chromobody GFP (AC-GFP) mRNA, large protrusions are colored green. Brackets indicated lamellipodia-like structure, arrows indicate filopodia branching out from dorsal tips of lamellipodia-like core. Asterisks indicate sclerotome cells, scale bar = 25 μm. (E, H) Summary of dorsal DM cell protrusions in (E) Wt and (H) *Da* embryos. Arrowheads indicate small protrusions, brackets indicate lamellipodia-like core structure of large protrusions, arrows indicate filopodia bundles branching off from dorsal tips of large protrusions. (I-I’’) Quantification of protrusions from Tg(*zic1:GFP*) (n = 14 embryos) and Tg(*zic1:GFP*);*Da* (n = 11 embryos). Protrusions of the 8^th^-12^th^ somite were counted (mean ± SD, *P_small protrusions_ = 0.01, ****P_large protrusions_ = 3.3e-08, ****P_total protrusions_= 2.1e-05). (J-K) Lateral view of protrusions extracted from time-lapse imaging of (A-B). Arrowheads indicate tip of protrusions, time is displayed in min. (J) Protrusion observed in Tg(*zic1:GFP*). (K) Protrusion observed in Tg(*zic1:GFP*);*Da*. Scale bar = 25 μm. Anterior = left, DM = dermomyotome, M = myotome, NT = neural tube.

To characterize the protrusions, we classified them according to their length into small (< 8 μm, Figure 3C arrowheads) and large (≥ 8 μm, Figure 3C arrow) protrusions (Figure 3 - figure supplement 1B-C). Based on their shape, we reasoned that the small protrusions correspond to lamellipodia (Figure 3C, arrowheads), while the large protrusions appeared more complex. To investigate the nature of large protrusions, we injected Actin-Chromobody GFP mRNA to visualize the actin skeleton. The large protrusions were found to exhibit a complex architecture consisting of a lamellipodia-like core structure (Figure 3D, bracket) with additional multiple bundles of filopodia (protrusions with linear arranged actin filaments) branching out from their dorsal tips (Figure 3D, arrows, summarized in Figure 3E).

Interestingly, time-lapse *in vivo* imaging of Tg(*zic1:GFP*);*Da* showed that in the *Da* background, protrusions started to form later (Figure 3B, Video 2) and the number of large protrusions and protrusions in total was significantly lower than in Wt (Figure 3I-I’’). In addition, protrusions were transient and mostly failed to form new branches at their dorsal tips (Figure 3K). While no difference in the actin skeleton of small protrusions could be observed, filament bundles branching out from large protrusions of *Da* DM cells contained fewer and shorter filopodia compared to Wt (Figure 3G, arrowheads, summarized in Figure 3H). These results indicate that the protrusive activity, especially the ability to form large protrusions, is significantly reduced in the *Da* mutant. The large protrusions of the dorsal DM cells continuously appeared at later stages of dorsal somite extension, too (Figure 3 - figure supplement 2).

Taken together, the unique large protrusions of the dorsal DM might be involved in guiding the epaxial myotome dorsally and *zic1* might promote this function.

Video 1: Onset of dorsal somite extension in Tg(*zic1:GFP*).

Dorsal view of time-lapse *in vivo* imaging of 24 ss Tg(*zic1:GFP*) embryo. 15^th^ somite is positioned in the center, z-stacks were imaged every 10 min, time is displayed in min.

Anterior = left, scale bar = 50 μm.

Video 2: Onset of dorsal somite extension in Tg(*zic1:GFP*);*Da*.

Dorsal view of time-lapse *in vivo* imaging of 24 ss Tg(*zic1:GFP*);*Da* embryo. 15^th^ somite is positioned in center, z-stacks were imaged every 10 min, time is displayed in min. Bright cell at the bottom migrating to the right is a melanophore. Anterior = left, scale bar = 50 μm.

### DM cells delaminate and accumulate between opposing somites during the late phase of dorsal somite extension

We continued to trace the behavior of the DM tip cells until the somites reach the top of the neural tube. Indeed, *in vivo* imaging of Tg(*zic1:GFP*) revealed that dorsal DM cells continued to form protrusions, and additionally, some of them delaminated to become Zic1-positive mesenchymal cells accumulating in the space between the dorsal ends of the left and the right somites (Figure 4A, star-shaped cells, arrowheads) from 4.5 dpf (stage 33) onwards. This is consistent with previous observations of strongly *zic1* expressing mesenchymal cells in Wt at late embryonic stages (Ohtsuka *et al*., 2004). As dorsal somite extension proceeded, the number of these mesenchymal cells increased, filling the space between the two myotomes (Figure 4A-D, arrowheads, Video 3-4, arrowheads indicate representative mesenchymal cells). These mesenchymal cells formed protrusions towards neighboring mesenchymal cells and DM cells at the tip of somites, creating a dense cellular network between the dorsal ends of the somites. Mosaic cell-labeling demonstrated that the mesenchymal cells originated from the DM (Figure 4 - figure supplement 1A-F, arrowheads). Interestingly, while mesenchymal cells dynamically formed protrusions, they showed no extensive migratory behavior and were rather stationary (Video 3-4, Figure 4 – figure supplement 2). This could suggest that these protrusions fulfill a non-migratory function. When the opposing somites came in contact with each other at 8 dpf (stage 38, one day before hatching), the mesenchymal cells tended to attach to the nearest DM cells at the tip of the somite, bridging the gap between the left and right DM cells (Figure 4K-K’’’’).

**Figure 4:**
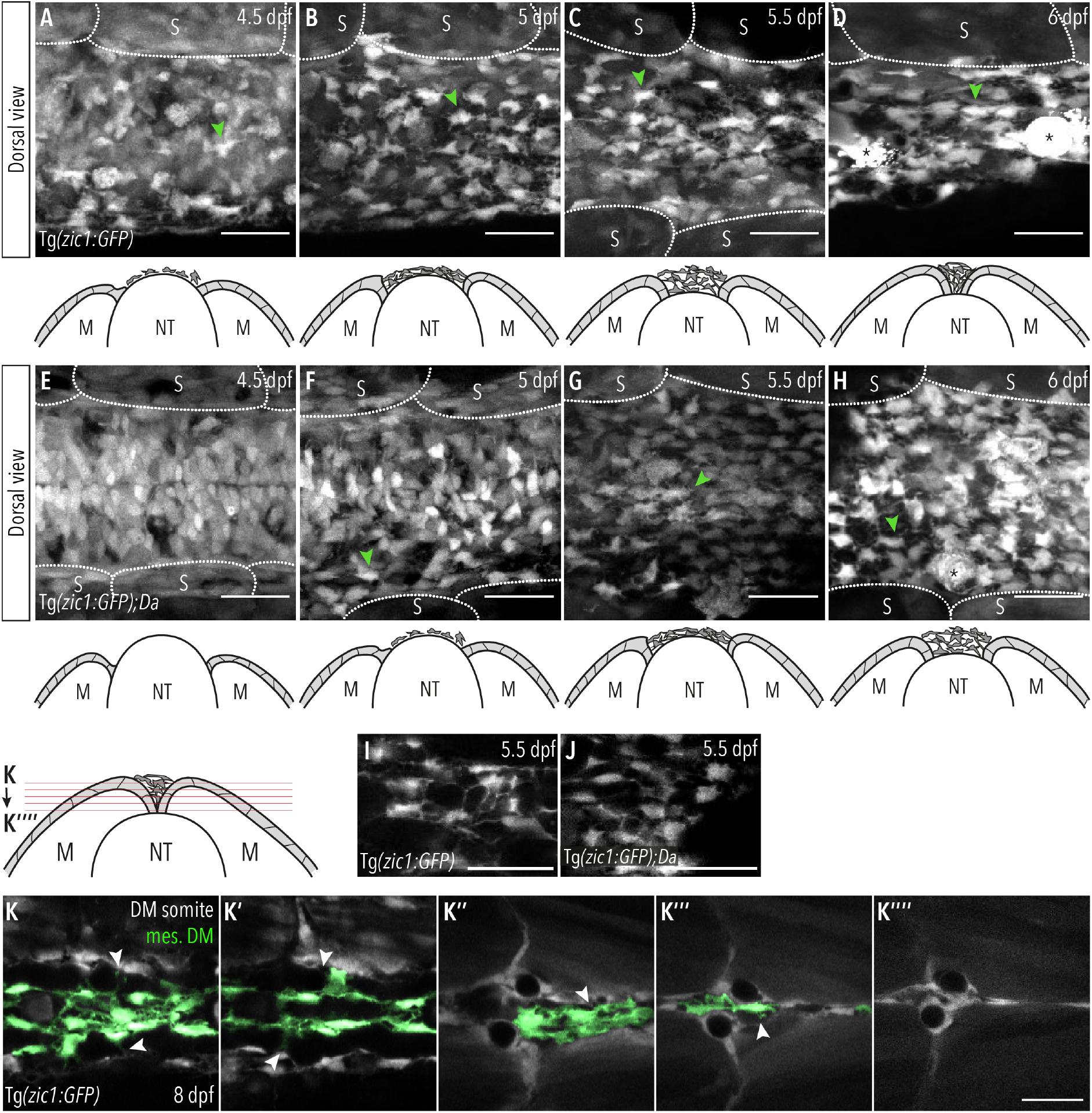
DM cells delaminate from the dorsal somite at the end of dorsal somite extension and accumulate between somites. (A-H) Dorsal view of maximum projection of Tg(*zic1:GFP*) at (A) 4.5 dpf, (B) 5 dpf, (C) 5.5 dpf and (D) 6 dpf and Tg(*zic1:GFP*);*Da* at (E) 4.5 dpf, (F) 5 dpf, (G) 5.5 dpf and (H) 6 dpf. 10^th^ somite positioned in center, arrowheads point to exemplary mesenchymal DM cells, asterisks mark melanophores. Corresponding schematic representation of cross-section of the trunk beneath each dorsal view. (I-J) Maximum projection of mesenchymal DM cells of Tg(*zic1:GFP*) (I) and Tg(*zic1:GFP*);*Da* (J). (K-K’’’’’) Consecutive z-planes of dorsal view of 10^th^ somite of Tg(*zic1:GFP*) embryo. Mesenchymal DM cells are colored green, arrowheads indicate protrusions formed between somitic DM cell and mesenchymal DM cell. Anterior = left, M = myotome, NT = neural tube, S = somite, scale bar = 25μm.

In *Da* mutants, mesenchymal DM cells were also detected in the space between the two myotomes, but the timing of their appearance was delayed, i.e. between 5 dpf (stage 34) and 5.5 dpf (stage 35) (Figure 4E-H) (4.5 dpf in Wt). Additionally, *Da* mesenchymal cells exhibited a rounder morphology and formed significantly fewer protrusions compared to Wt (Figure 4I-J, Figure 4 - figure supplement 1G-L).

Collectively, dorsal DM cells and the mesenchymal cells derived from them seem to actively participate in the entire process of dorsal somite extension, from its onset to neural tube coverage at the end.

Video 3: Mesenchymal DM cells during dorsal somite extension at 4.5 dpf.

Dorsal view of time-lapse *in vivo* imaging of 4.5 dpf Tg(*zic1:GFP*) embryo. 10^th^ somite positioned in center, z-stacks were imaged every 10 min, time is displayed in min.

Arrowhead indicates representative mesenchymal DM cell. Anterior = left, scale bar = 50 μm.

Video 4: Mesenchymal DM cells during dorsal somite extension at 5.5 dpf.

Dorsal view of time-lapse *in vivo* imaging of 5.5 dpf Tg(*zic1:GFP*) embryo. 10^th^ somite is positioned in center, z-stacks were imaged every 10 min, time is displayed in min.

Arrowhead indicates representative mesenchymal DM cell. Anterior = left, scale bar = 50 μm.

### Zic1 regulates the expression of dorsal-specific genes during somite differentiation

We then addressed the molecular machinery controlling the dorsal somite extension investigated above. Since somite extension is impaired in the *zic1*-enhancer mutant *Da* (Moriyama *et al*., 2012; Kawanishi *et al*., 2013), we reasoned that downstream genes of Zic1 are regulators of this process.

First, we identified genes which are specifically expressed in the dorsal somites. For this, we dissected somites of the transgenic line Tg(*zic1:GFP*) and FACS sorted them into cells from the dorsal (GFP+) and ventral (GFP-) somites, including the DM, and performed RNA-seq and ATAC-seq on both cell populations (Figure 5A, Figure 5 - figure supplement 1A). The RNA-seq identified 1,418 differentially expressed genes. Among them 694 genes showed higher expression in the dorsal somites (termed hereafter dorsal-high genes), and 724 genes showed higher expression in the ventral somites (termed hereafter dorsal-low genes) (Figure 5C). We confirmed that *zic1* and *zic4* were found among the dorsal-high genes (Figure 5C, Supplementary table 1).

**Figure 5:**
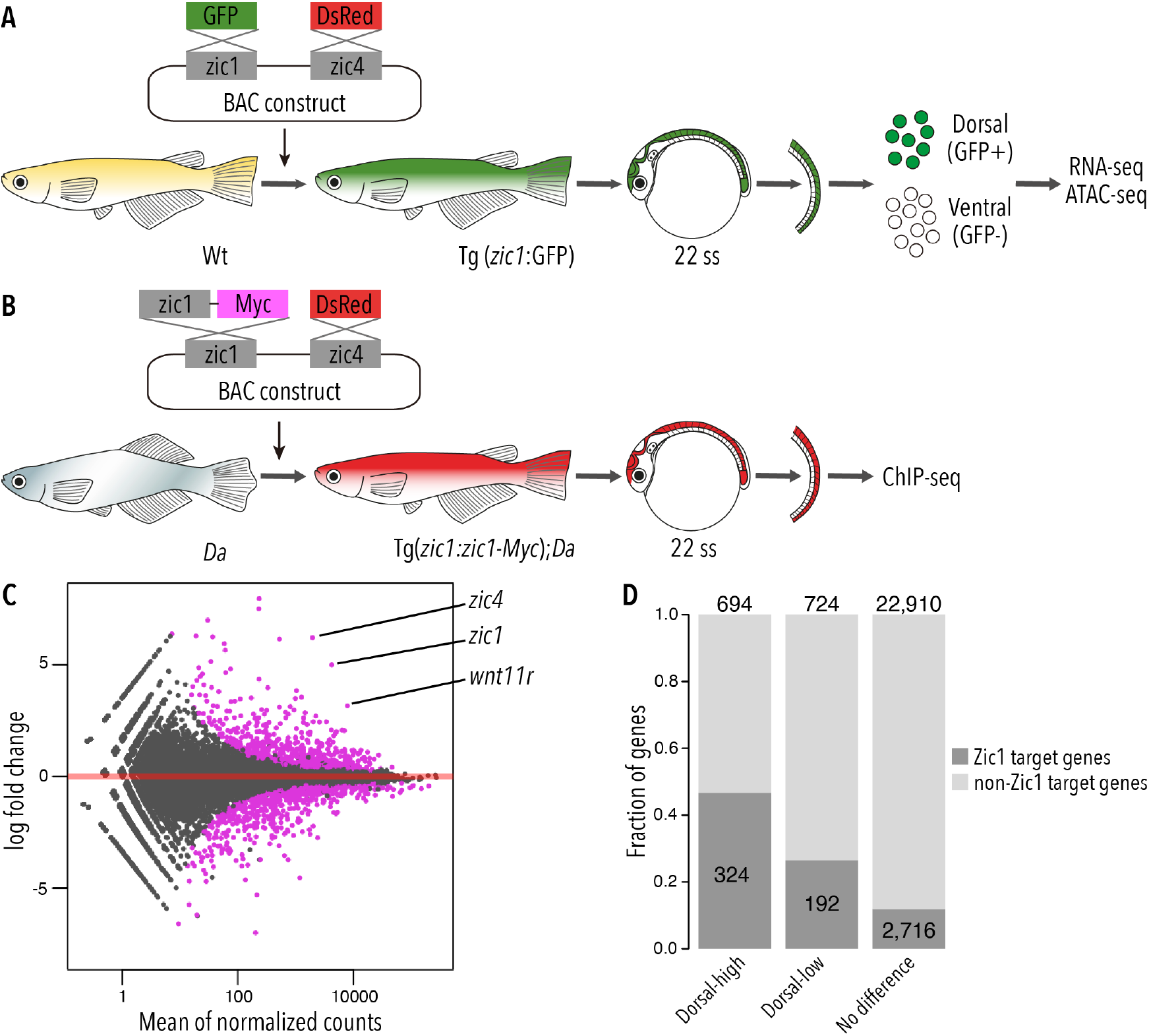
Zic1 regulates dorsal-specific expression of genes in the somites. (A) Schematic representation of preparation of dorsal and ventral somite cells for RNA-seq and ATAC-seq. (B) Schematic representation of generating the transgenic line Tg(zic1:*zic1-Myc*)*;Da.* Somites expressing *zic1-Myc* are subjects to ChIPmentation against Myc. (C) Analysis of RNA-seq revealed that 694 genes are expressed specifically in the dorsal somites (among these genes are *zic1* and *zic4*) and 724 genes are expressed specifically in ventral somites. Magenta indicates differentially expressed genes (adjusted p-value<0.01). (D) The ChIPmentation against Zic1 revealed that 324 of dorsal-high genes, 192 dorsal-low genes and 2,716 non differential expressed genes in the somites are potential Zic1 target genes.

Next, we identified potential direct Zic1-target genes by investigating Zic1 binding sites. Since there were no suitable antibodies available to perform ChIP-seq against medaka Zic1, we created a transgenic line expressing a Myc tagged *zic1* in the *Da* background under the control of *zic1* promoter and enhancers (Tg(*zic1*:*zic1-Myc;zic4:DsRed*);*Da,* called *Tg(zic1:zic1-Myc);Da* hereafter). This transgene *(zic1:zic1-Myc)* efficiently rescued the ventralized phenotype (dorsal and anal fin shape, pigmentation pattern, and body shape) of the *Da* mutant (Figure 5 - figure supplement 1B-C), verifying the full functionality of the tagged Zic1 protein. Somites of the transgenic line *Tg(zic1:zic1-Myc);Da* were dissected and subjected to ChIP-seq using antibodies against Myc (Figure 5B). Since somites contain a low number of cells, we applied ChIPmentation (Schmidl *et al*., 2015) to identify genome-wide Zic1 binding sites. From two biological replicates, 5,247 reliable ChIP peaks were identified, and we confirmed that the enriched DNA motif among these peaks was showing high similarity with previously identified binding motifs of ZIC family proteins (Figure 5 - figure supplement 2A-B). Then, we associated each Zic1 peak to the nearest gene within 50 kb and identified 3,232 genes as Zic1 target genes. By comparing the ChIPmentation with RNA-seq data, we found that Zic1 target genes are overrepresented in dorsal-high and dorsal-low genes. While 47% of the dorsal-high genes and 27% of the dorsal-low genes were found to be Zic1 target genes, only 12% of genes which showed no differential expression in dorsal or ventral somites were potential Zic1 downstream target genes (Figure 5D, Supplementary table 1). This suggests that Zic1 can function as transcriptional activator and repressor of versatile genes, but the former role seems to be dominant.

### *Wnt11r* is a direct downstream target of Zic1 and down-regulated in the dorsal somites of the *Da* mutant

To identify potential regulators of dorsal somite extension, we further investigated the differentially expressed Zic1 target genes. Gene Ontology (GO) analysis indicated that both dorsal-high and dorsal-low gene groups, regardless of whether they are Zic1 targets or non-Zic1 targets, were significantly enriched in development related GO terms (Figure 6A, Figure 6 - figure supplement 1A, Supplementary table 2, 3). This indicates that Zic1 regulates a number of developmental genes both directly and indirectly. These results are consistent with the fact that Zic1 regulates various dorsal-specific morphologies of somite-derivatives (Kawanishi *et al*., 2013).

**Figure 6:**
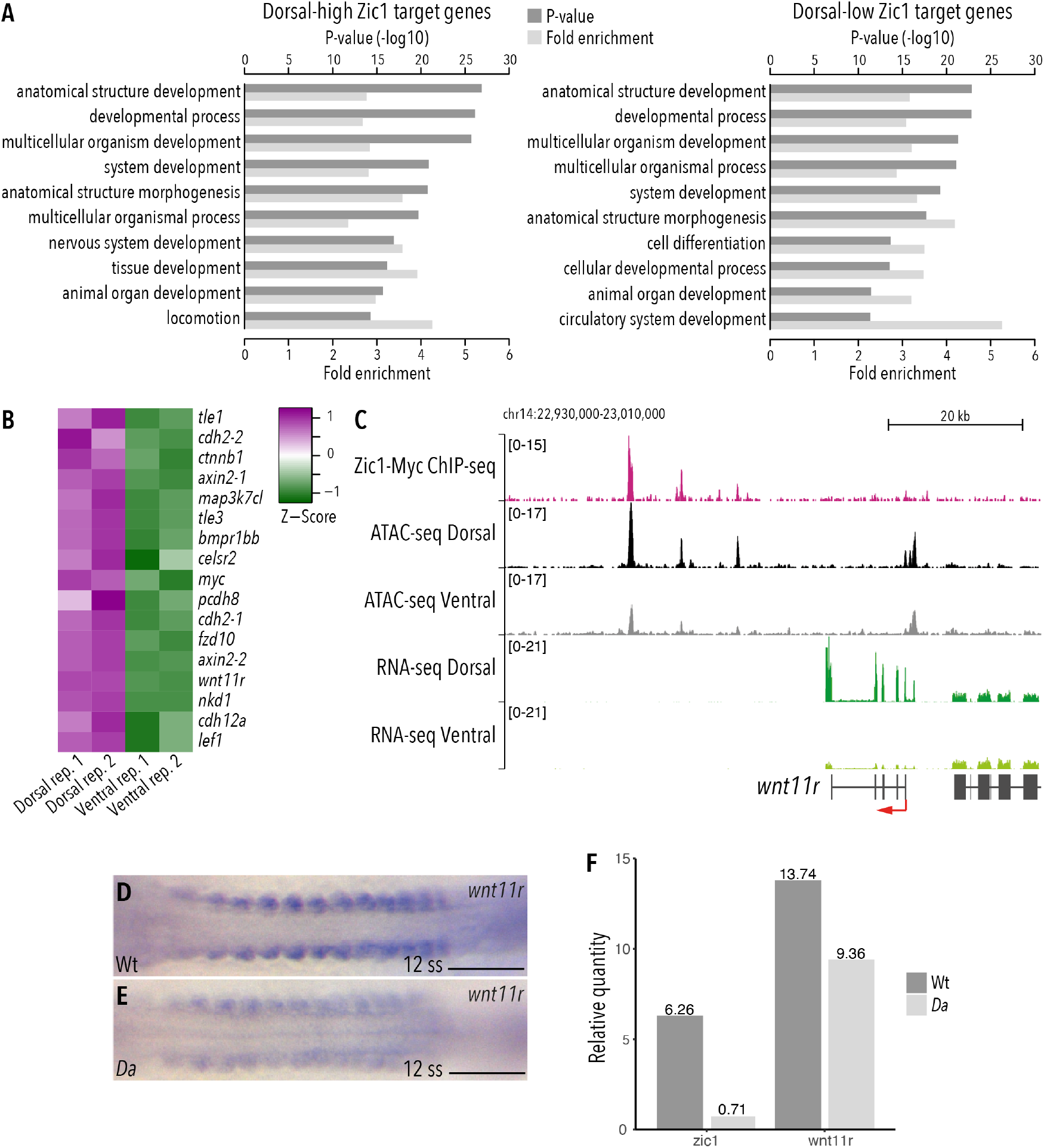
*Wnt11r* is a direct downstream target of Zic1 and down-regulated in the dorsal somites of the *Da* mutant. (A-A’) GO analysis of (A) dorsal-high and (A’) dorsal-low Zic1 target genes. (B) Pathway enrichment analysis indicated that genes associated with Wnt signaling pathway are specifically enriched in dorsal-high Zic1 target genes. (C) Analysis of the *wnt11r* locus. Peaks form the ChIPmentation against Zic1 (magenta track) overlap with open chromatin regions in the dorsal somites (black track), while this genomic region is less open in ventral somites (grey track). RNA-seq data revealed that *wnt11r* is highly expressed in dorsal somites (dark green track) and minimally in ventral somites (light green track). (D-E) Dorsal view of tails of whole-mount *in situ* hybridization against *wnt11r* performed in (D) Wt and (E) *Da* 12 ss embryos. The expression of *wnt11r* is reduced in the *Da* mutant. (F) RT-PCR performed on pooled tails of Wt and *Da* 12 ss embryos indicates that *zic1* expression is reduced by 8.8-fold and *wnt11r* expression by 1.4-fold in the *Da* mutant. Anterior = left, scale bar = 100 μm.

In the dorsal-high Zic1 targets, GO terms related to cell migration showed higher enrichment (e.g. “chemotaxis” (P=5.62E-10), “locomotion” (6.65E-15), “ameboidal-type cell migration” (P=9.32E-10)) than dorsal-low genes or non-Zic1 target genes (Supplementary table 3). Interestingly, Wnt signaling pathway genes (e.g.: *axin2, wnt11r, sp5, lrp5, fzd10, prickle1a*) and semaphorin-plexin signaling pathway genes (e.g.: *sema3a, sema3c, sema3g, plxna1, plxna2, plxnb2, plxnb3, nrp2*) were included in these gene groups. Indeed, the GO terms “Wnt signaling pathway” and “Semaphorin-plexin signaling pathway” themselves were also significantly enriched (P=4.45E-7 and 1.94E-8, respectively) in dorsal-high Zic1 target genes (Supplementary table 2). This suggests that Zic1 directly regulates Wnt pathway, semaphorin and plexin genes, possibly to regulate cell movement in the dorsal somites. We also noticed that the terms “extracellular matrix organization” (P=7.74E-7; e.g.: *adamts20, fbln1*), “cell communication” (P=1.95E-5; e.g.: *efna5, epha3*) are enriched, suggesting that these genes also affect dorsal somite cell behavior.

Wnt signaling pathway components also ranked high among the dorsal-high Zic1 target genes by pathway enrichment analysis (Figure 6B, Figure 6 - figure supplement 1B). Among them, we focused on *wnt11r* for further analyses due to the following reasons: First, *wnt11r* was one of the most differentially expressed genes in dorsal somites (Figure 5C, Supplementary table 1). Second, previous studies implicated Wnt11 in protrusion formation and cell migration (Ulrich *et al*., 2003; De Calisto *et al*., 2005; Garriock and Krieg, 2007; Matthews *et al*., 2008). This is particularly interesting since DM cells also exhibit protrusions and migration activity during dorsal somite extension, which are defective in *Da* mutants.

At the *wnt11r* locus, peaks of the Zic1-ChIP overlapped with intergenic open chromatin regions downstream of *wnt11r*. These sites were more accessible in dorsal somites than in ventral somites, suggesting that Zic1 regulates *wnt11r* via enhancers (Figure 6C).

Additionally, *in situ* hybridizations against *wnt11r* performed on Wt and *Da* embryos indicated that *wnt11r* expression is significantly reduced at the dorsal tip of *Da* dorsal somites including the DM (Figure 6 - figure supplement 1C-D). Already when the *zic1* expression becomes restricted in the dorsal somites (12 ss) and with proceeding development when *zic1* expression gets further restricted to the most dorsal part of the somites and mesenchymal DM cells (Ohtsuka *et al*., 2004), the expression of *wnt11r* in the dorsal somites of the *Da* mutant was reduced compared to Wt (Figure 6D-F, Figure 6 - figure supplement 1C-F’).

Taken together, we identified *wnt11r* as promising downstream target gene of Zic1, which could be a novel somite dorsalization factor and play a role in dorsal somite extension.

### Wnt11r regulates cell behavior of dorsal DM

Our previous RNA-seq dataset revealed that the expression of *wnt11r* starts before gastrulation and increases as development proceeds (Figure 7 - figure supplement 1A)(Nakamura *et al*., 2021). This makes it challenging to examine the role of *wnt11r* during late embryonic development using loss-of-function experiments. We took two different approaches to knock-down Wnt11r during dorsal somite extension. Firstly, we used a *wnt11r* anti-sense morpholino (*wnt11r* MO) and determined a concentration which resulted in maximal knock-down effects during dorsal somite extension with minimal gastrulation phenotypes (we excluded any embryos showing gastrulation defects in our experiments). Secondly, we performed temporally controlled knock-down of Wnt11r after gastrulation using photo-cleavable Photo-Morpholinos (PhotoMOs) (Tallafuss *et al*., 2012) (Figure 7A). Strikingly, in *wnt11r* Photo-Morphants with severe phenotypes the dorsal myotome failed to cover the neural tube (n = 7, Figure 7B-C), a phenotype similar to the *Da* mutant myotome. Additionally, the myofibers of the Photo-Morphants at 9 dpf were shorter and less organized. This phenotype is consistent with a previously reported role of Wnt11 during early myogenesis, where it regulates the elongation and orientation of myoblasts (Gros, Serralbo and Marcelle, 2008).

**Figure 7:**
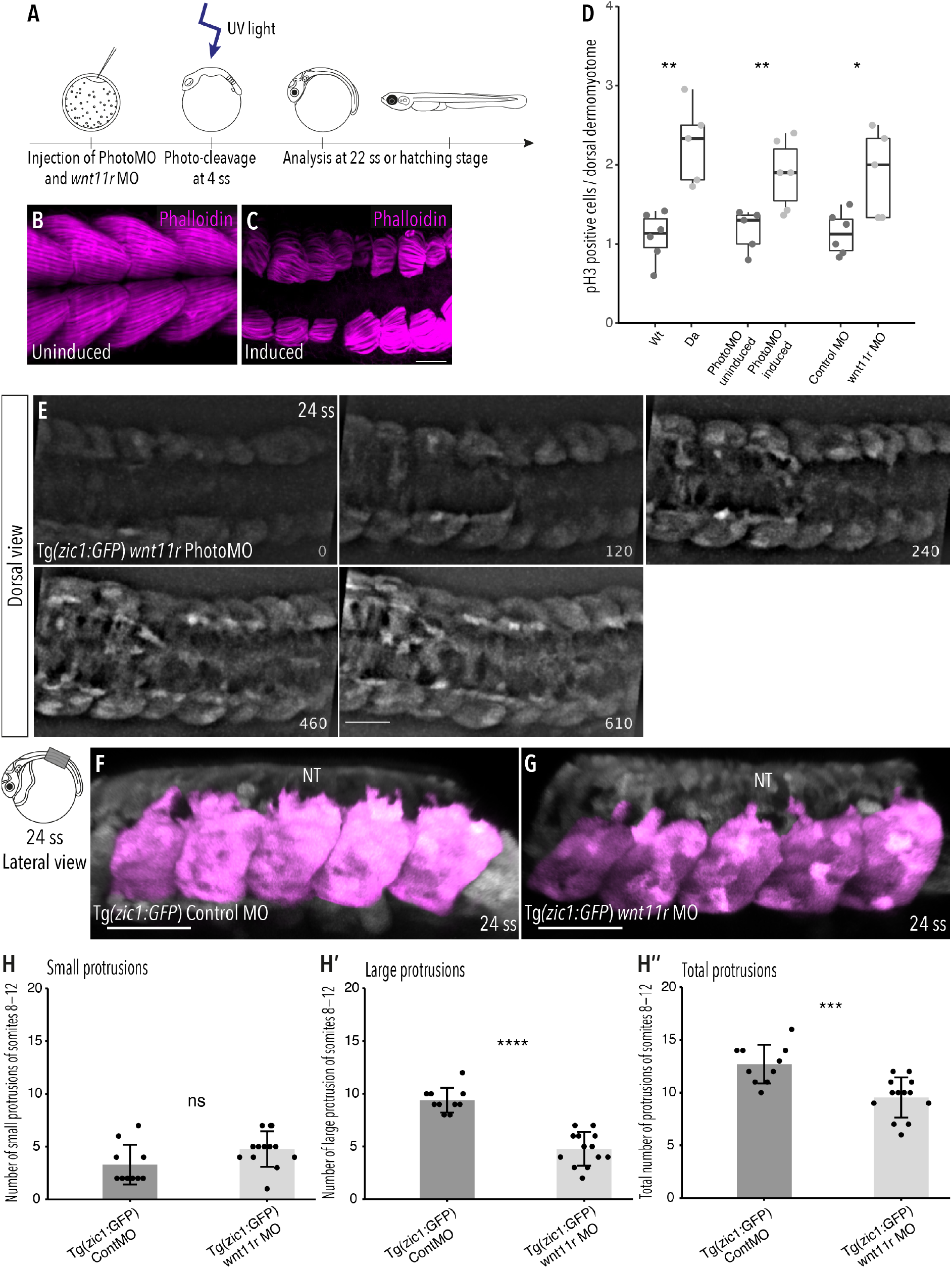
Knock-down of Wnt11r in Wt embryos recapitulates the *Da* dorsal somite phenotype. (A) Schematic outline of PhotoMO mediated knock-down of Wnt11r. (B,C) Dorsal view of maximum projection of whole-mount Phalloidin staining of 9 dpf embryos injected with (B) PhotoMO but not photocleaved and (C) photo-cleaved. The epaxial myotomes of the *wnt11r* morphant does not cover the neural tube. (D) Quantification of PH3-positive cells in the dorsal DM of 22 ss embryos. Embryos with reduced levels of Wnt11r in their dorsal somites (n = 40.5 somites from 5 *Da* embryos, n = 61.5 somites of 6 *wnt11r* PhotoMO-Morphant embryos and n = 52 somites from 5 *wnt11r* morphant embryos) have significantly more PH3-positive cells in their dorsal DM compared to the respective controls (n = 66 somites from 6 Wt embryos, n = 51 somites of 5 uninduced PhotoMO embryos and n = 65 somites from 6 Control MO embryos) (median, first and third quartiles, **P_Wt/*Da*_ = 0.0035, **P_PhotoMO_ = 0.0091, *P_MO_ = 0.031). (E) Dorsal view of time-lapse imaging of 24 ss Tg(*zic1:GFP*) embryo injected with PhotoMO-douplex and photo-cleaved at 4 ss. Z-stacks were recorded every 10min, time is displayed in min, 15^th^ somite is positioned in the center. (F-G) Lateral view of 3D-reconstructions of dorsal somites (magenta) of Tg(*zic1:GFP*) injected with (F) Control MO and (G) *wnt11r* MO. (H-H’’) Quantification of protrusions formed by the 8^th^-12^th^ somite of Tg(*zic1:GFP*) injected with Control MO (n = 10 embryos) or *wnt11r* MO (n = 13 embryos) (mean ± SD, ^ns^P_small protrusions_ = 0.069, ****P_large protrusions_ = 7.7e-08, ***P_total protrusions_= 0.00064). Anterior = left, NT = neural tube, scale bar = 50 μm.

Next, we investigated the proliferative activity of dorsal DM cells in *wnt11r* morphants and found that knock-down of *wnt11r* (either by conventional MO or by PhotoMO) induced significantly more PH3-positive cells per somite, compared to control embryos at 22 ss (Figure 7D). These findings are similar to the observations previously made in the dorsal DM of *Da* mutants, and suggest that Wnt11r is negatively regulating proliferation of dorsal DM cells.

We then explored whether the onset of dorsal somite extension in *wnt11r* morphants is similarly impaired as in the *Da* by time-lapse *in vivo* imaging. Remarkably, we observed a protrusion formation behavior of *wnt11r* morphant DM cells similar to *Da* DM cells, namely delayed onset of protrusion formation and the formation of fewer, shorter protrusions (Figure 7E, Video 5). Quantification of protrusions indicated that *wnt11r* morphants have significantly fewer large protrusions and protrusions in total, compared to control embryos (Figure 7F-H’’).

Overall, the knock-down of the Zic1 target gene Wnt11r recapitulated the phenotype of *Da* DM cells (Figure 3G-G’’), showing the essential role of Wnt11r in regulating protrusion formation of DM cells.

Video 5: Onset of dorsal somite extension in *wnt11r* morphant embryo.

Wnt11r was knocked-down using the PhotoMO approach (Figure 7A) in a Tg(*zic1:GFP*) embryo. Dorsal view of time-lapse *in vivo* imaging of 24 ss Tg(*zic1:GFP*) embryo. 15^th^ somite is positioned in the center, z-stacks were imaged every 10 min, time is displayed in min. Anterior = left, scale bar = 50 μm.

To further confirm the importance of Wnt11r during dorsal somite extension, we performed rescue experiments in *Da* embryos. At 18 ss (2.1 dpf, stage 25), we injected a mix of human recombinant Wnt11 (hrWnt11) protein (or BSA in the control group) and Dextran Rhodamine onto the top of the 10^th^ somite of Tg*(zic1:GFP);Da* embryos (Figure 8A, B). Strikingly, the dorsal DM of *Da* embryos injected with Wnt11 formed significantly more large protrusions and more protrusions in total compared to *Da* embryos injected with BSA only (Figure 8C-C’’). This indicates that the ventralized *Da* DM protrusion phenotype can be partially rescued by Wnt11 protein injections and further emphasizes the importance of Wnt11r in somite dorsalization and during dorsal somite extension.

**Figure 8:**
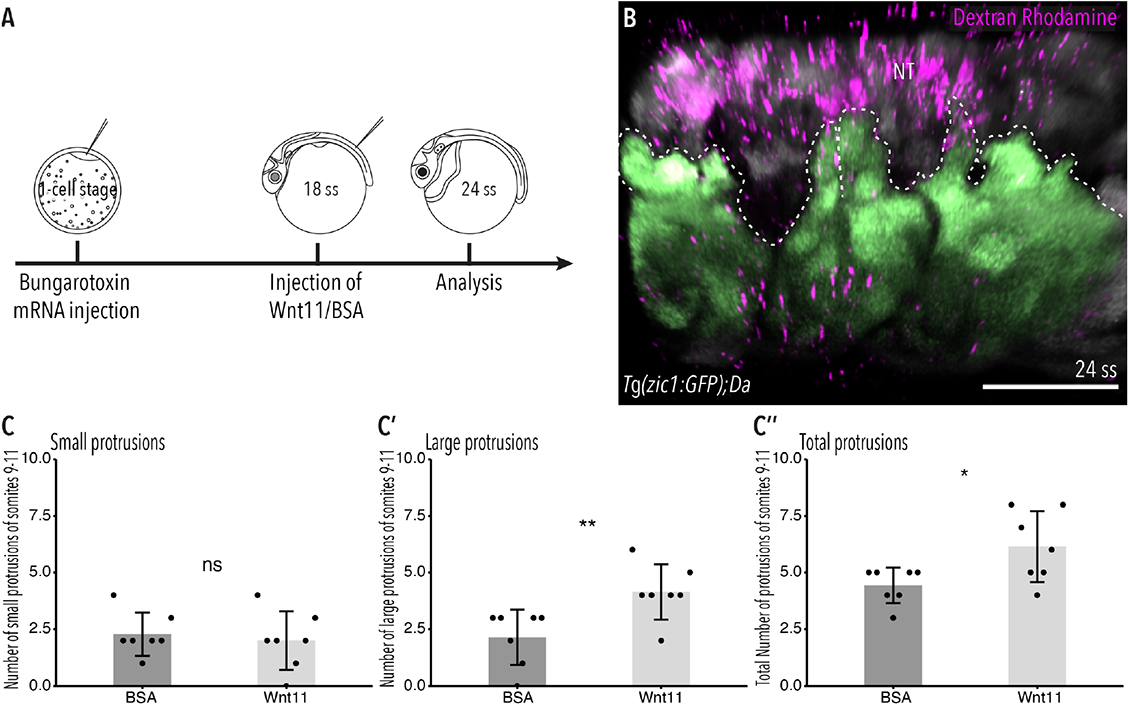
Wnt11 injections partially rescue the Tg(*zic1:GFP*);*Da* protrusion phenotype. (A) Schematic representation of Wnt11 protein injection on top of the 10^th^ somite of 18 ss Tg(*zic1:GFP*);*Da)* embryos. (B) Dorsal view of 3D-reconstruction of *in vivo* imaging of dorsal somites (green) of Tg(*zic1:GFP*);*Da* embryos injected with Wnt11r-Dextran Rhodamine mix (magenta). Protrusions are outlined. (C-C’’) Quantification of dorsal protrusions of the 9^th^ – 11^th^ somite (somite at injection site plus adjacent somites) of embryos injected with BSA (n = 7 embryos) or hrWnt11(n = 7 embryos) (mean ± SD, ^ns^P_small protrusions_ = 0.65, **P_large protrusions_ = 0.0095, *P_total protrusions_ = 0.03). Anterior = left, dorsal = up, NT = neural tube, scale bar = 50 μm.

### Wnt11r acts through the Wnt/Ca^2+^ signaling pathway at the onset of somite extension

Finally, we investigated through which signaling pathway the non-canonical Wnt11r acts. In *Xenopus*, Wnt11r acts through the Wnt/Ca^2+^ pathway regulating the migration of cells from the dorsal somite and the neural crest into the dorsal fin fold (Garriock and Krieg, 2007). To examine whether this signaling pathway also plays a role during dorsal somite extension, we inhibited the Wnt/Ca^2+^ signaling pathway using KN-93 in Tg(*zic1:GFP*) embryos from 4 ss (1.3 dpf, stage 20) to 22 ss (Figure 9A). KN-93 is known to inhibit CaMKII, a component of the Wnt/Ca^2+^ pathway (Wu and Cline, 1998; Garriock and Krieg, 2007; Rothschild *et al*., 2013). Embryos treated with KN-93 showed a higher number of PH3-positive dorsal DM cells per somite, compared to embryos in the control group (Figure 9B). Furthermore, DM cells at the tip of the dorsal somite of embryos treated with KN-93 formed significantly fewer large and fewer protrusions in total, compared to embryos in the control group (Figure 9C-C’’), although the effect was less significant compared to the *wnt11r* morphants (Figure 7).

**Figure 9:**
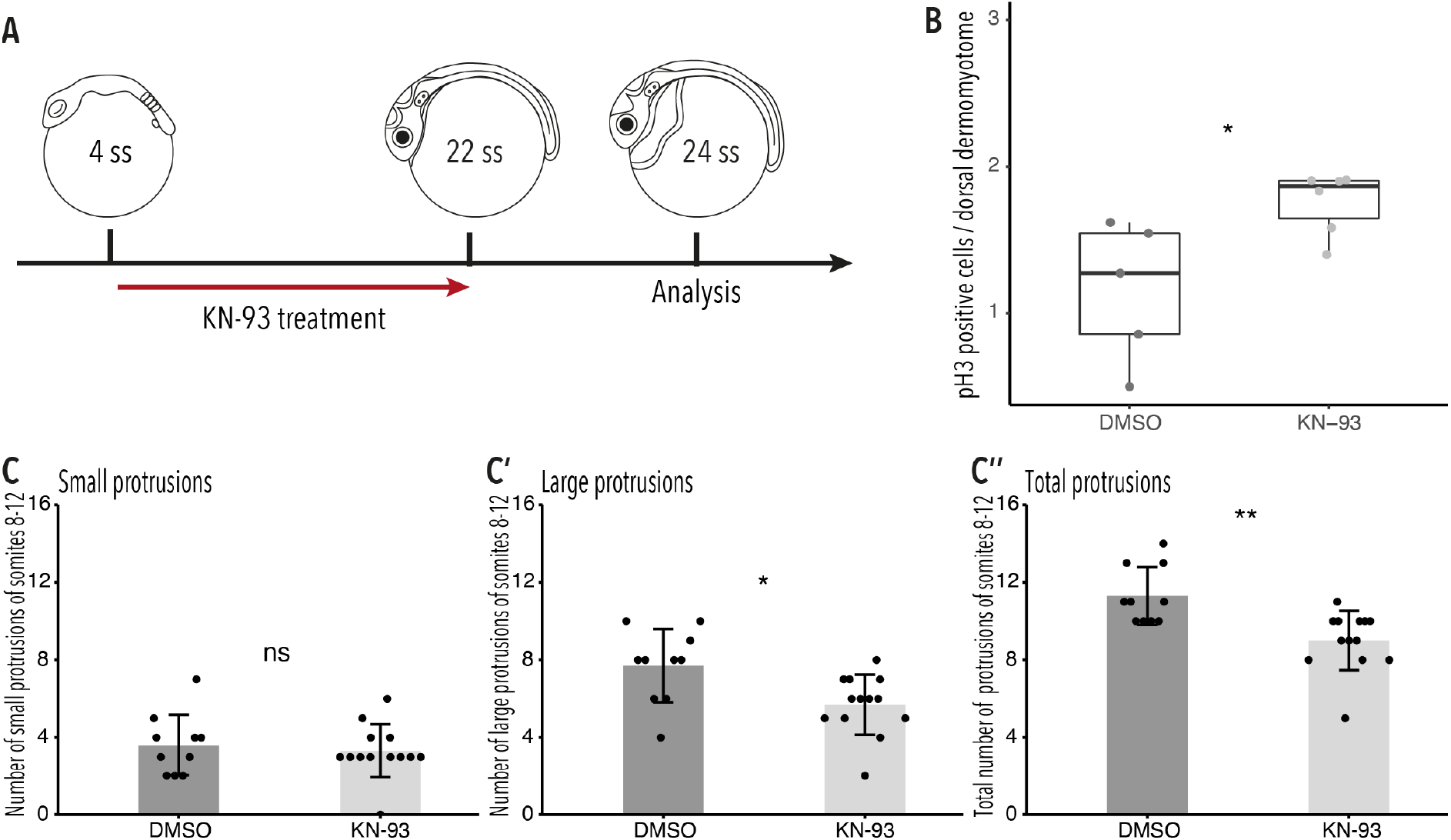
Wnt11r acts through the Wnt/Ca^2+^ signaling pathway during dorsal somite extension. (A) Schematic representation of KN-93 treatment, embryos in the control group were treated with DMSO. (B) Embryos treated with KN-93 (n = 65.5 somites from 6 embryos) have significantly more PH3-positive cell in the dorsal DM compared to embryos of the control group (n = 52.5 somites from 5 embryos) (median, first and third quartiles, *P = 0.045). (C-C’’) Quantification of protrusions formed by the 8^th^-12^th^ somites of Tg(*zic1:GFP*) embryos treated with DMSO (n = 10 embryos) or KN-93 (n = 13 embryos)(mean ± SD, ^ns^P_small protrusions_ = 0.65, *P_large protrusions_ = 0.014, **P_total protrusions_ = 0.0017).

From these results, we suggested that Wnt11r potentially acts through the Wnt/Ca^2+^ signaling pathway at the onset of somite extension. Wnt/Ca^2+^ signaling pathway is known to regulate actin polymerization (Choi and Han, 2002; Kohn and Moon, 2005) which could explain the dynamic protrusive activity of the DM cells (Figure 3), emphasizing the importance of Wnt11r during epaxial myotome morphogenesis.

## Discussion

Here, we elucidate the key developmental process underlying the epaxial myotome morphogenesis in teleost fish, medaka. At the initiation of dorsal somite extension, DM cells at the tip of the dorsal somite form unique large, motile protrusions, which extend dorsally, and potentially play a pioneering role in guiding the myotome towards the top of the neural tube. By analyzing dorsal somite extension in the ventralized *Da* mutant, we demonstrated that *zic1* is essential for this process. DM cells of the *Da* dorsal somite have a higher proliferative activity and form fewer and shorter protrusions during dorsal somite extension. These altered cellular properties of *Da* dorsal DM cells, together with a delayed onset of dorsal somite extension, potentially cause the incomplete coverage of the neural tube by the epaxial myotome at the end of embryonic development.

Furthermore, we investigated the molecular background of dorsal somite extension and identified a direct downstream target of Zic1, the non-canonical Wnt *wnt11r*, as a crucial factor for dorsal somite extension. Wnt11r reduces the proliferative activity of DM cells in the dorsal somites, but instead, promotes the formation of large, motile protrusions of dorsal DM cells at the tip of the somite. Indeed, *wnt11r* morphants recapitulate the phenotype of the *Da* mutant. Additionally, the protrusion phenotype of *Da* dorsal DM cells can be partially rescue by injection of Wnt11 proteins. Based on these findings, we propose a model for dorsal somite extension shown in Figure 10.

**Figure 10:**
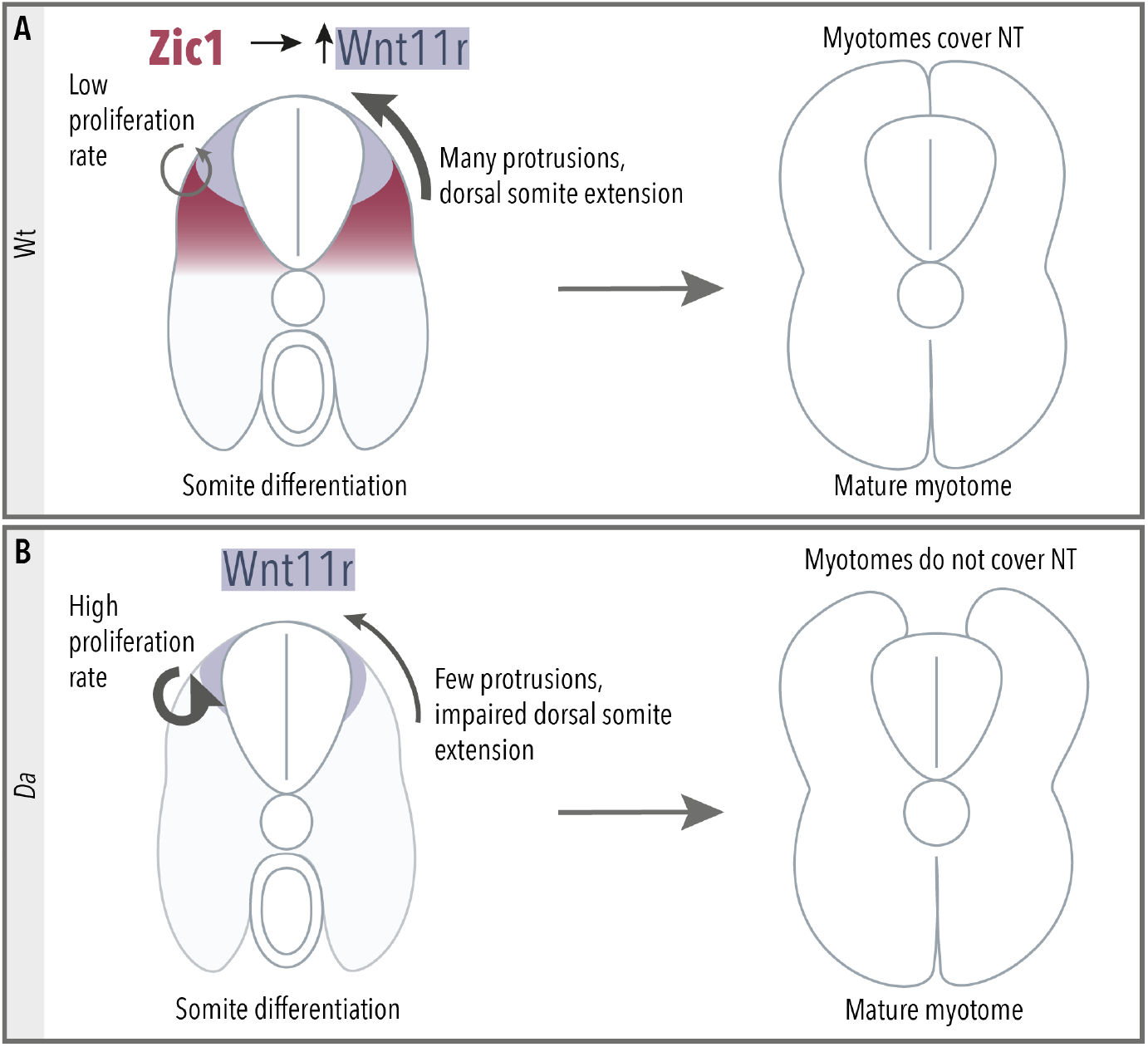
Summary of dorsal somite extension in Wt and the *Da* mutant. (A) Zic1 induces or maintains the expression of *wnt11r* during somite differentiation. This leads to a reduced proliferative activity of the dorsal DM and increases protrusion formation of the dorsal DM cells. Ultimately, the DM guides the epaxial myotomes dorsally where the myotomes form a gapless muscle layer covering the neural tube at the end of embryonic development. (B) In the *Da* mutant *wnt11r* expression is reduced in the dorsal somites. The dorsal DM cells show a high proliferative activity and reduced ability to form numerous large protrusions causing insufficient dorsal somite extension. This results in the incomplete coverage of the neural tube by the epaxial myotomes at the end of embryonic development.

In the present study, we described the characteristic behavior of dorsal DM cells and suggest their guiding role for the epaxial myotome moving towards the top of the neural tube. During dorsal somite extension, DM cells at the tip of the dorsal somites form large, motile protrusions which contain multiple bundles of filipodia-like protrusions dynamically branching out from the tip of the protrusions. These long protrusions could be beneficial for invading the restricted open space between the neural tube and the ectoderm. Additionally, previous studies showed that in migrating mesenchymal cells, the formation of lamellipodia is associated with higher migratory speed, whereas filopodia play an exploratory role and are associated with high directionality (Leithner *et al*., 2016; Innocenti, 2018). Large protrusions of wild-type DM cells consist of a lamellipodia-like core and multiple filopodia, which could account for fast dorsal somite extension with high accuracy. The detailed structure and function of the large protrusions needs to be further investigated in the future studies.

At later stages of dorsal somite extension, we observed that DM cells delaminate from the tip of the dorsal somites and progressively occupy the space between the opposing dorsal somites. These mesenchymal DM cells actively form protrusions towards neighboring mesenchymal DM cells and DM cells of the somites, thus forming a dense cellular network on top of the neural tube. This could provide a communication platform for the opposing somites to meet at the right position, exactly on top of the neural tube.

The ability of the motile DM cells to guide the underlying myotome dorsally assures complete coverage of the neural tube by the epaxial myotomes. Considering the fact that DM cells are in close cell-cell contact with the underlying myotome, it is plausible that they are biomechanically coupled with myotomal cells, facilitating dorsal myotome extension. A similar mechanism has been investigated in the mouse neural tube formation. During mouse neural tube closure, the surface ectoderm, overlying the neuroepithelium, forms cell protrusions towards the midline. Disruption of membrane ruffles, a form of lamellipodia, in the surface ectoderm results in incomplete fusion of the subjacent neuroepithelium (Rolo *et al*., 2016).

In medaka dorsal somites, Wnt11r exerts its effect through promotion of protrusion formation and down-regulation of cell proliferation in the dorsal DM. Regarding the protrusion formation, previous studies of the zebrafish mutant *silberblick* and migrating neural crest cells consistently reported that Wnt11 is involved in the oriented elongation and stabilization of protrusions (Ulrich, 2003; De Calisto *et al*., 2005; Matthews *et al*., 2008). Similarly, myocardial cells in the heart of *Wnt11* mutant mice also form fewer protrusions (Zhou *et al*., 2007). Thus, regulation of cell protrusions is a conserved function of Wnt11, observed in various developmental processes. However, the relationship between Wnt11 and cell proliferation could be context-dependent, as it negatively regulates proliferation in mouse neonatal hearts (Touma *et al*., 2017), while it promotes cell proliferation in mouse intestinal epithelial cell culture (Ouko *et al*., 2004). Hence, the function of Wnt11r in medaka somites is unique in that it promotes tissue elongation by regulating a balance between proliferative and migrative activity.

By inhibiting the Wnt/Ca^2+^ signaling pathway we showed that Wnt11r probably acts through this non-canonical Wnt signaling pathway during dorsal somite extension. Likewise, previous studies in *Xenopus* have shown that Wnt11r acts through this pathway during cell migration into the dorsal fin fold. There it regulates epithelial–mesenchymal transition in a distinct dorsal somite cell population which, together with a population of neural crest cells, contribute to the mesenchyme of the dorsal fin fold (Garriock and Krieg, 2007). Furthermore, during convergent extension in vertebrate gastrulation, Wnt/Ca2^+^ pathway can regulate cell adhesion by promoting actin polymerization (Choi and Han, 2002; Kohn and Moon, 2005).

Importantly, our RNA-seq and ChIPmentation analyses revealed that Zic1 has diverse downstream target genes including various developmental genes. This suggests that the dorsal myotome is established via pleiotropic actions of Zic1; Wnt11r may not be a sole factor for dorsal somite extension, although it was shown to be essential in the present study. The semaphoring-plexin pathway may play a role, since previous studies suggested, besides its implication in axon guidance and neural cell migration, a role in non-neural cell migration (Alto and Terman, 2017). Furthermore, cell migration and tissue deformation are often linked with the extracellular matrix (ECM), which provides guiding or restraining cues influencing cell movements. Previous studies showed that during mouse epaxial myotome development, ECM composition changes dynamically, which is tightly accompanied by epaxial muscle morphogenesis; while the laminin content in the ECM decreases, increasing tenascin and stable fibronectin contents potentially promotes the alignment of myofibers and their final organization (Deries *et al*., 2012). Furthermore, recent studies suggested that cells can remodel the surrounding ECM and thereby increase their own motility. For example, during zebrafish gastrulation, the metalloproteinase *mmp14* is expressed by migrating endoderm cells and degrades laminin and fibronectin, components of the ECM (Hu et al., 2018). In this context, of particular importance is our identification of *Adamts20*, encoding a proteoglycanase, as a dorsal-high Zic1-target gene in the somites (Supplementary table 1). Since Adamts20 is known to play a pivotal role in embryonic melanoblast migration by remodeling the dermal ECM (Rao *et al*., 2003; Silver *et al*., 2008), Adamnts20 could thus facilitate migration of Wnt11r-expressing DM cells through remodeling of the ECM. Further functional analysis of dorsal-high Zic1-target genes identified in somites will provide useful insight into the molecular network driving dorsal somite extension.

Finally, in vertebrates and especially in fish, body shape and muscle morphology are closely linked, since a majority of the body mass consist of muscular tissue. Previous studies in fish populations have shown that speciation and adaptation to a specific aquatic habitat are associated with a changes in body depth, a measurement of the trunk dorsoventral axis (Tobler *et al*., 2008; Elmer *et al*., 2010; Weese, Ferguson and Robinson, 2012; Fruciano *et al*., 2016). In this context, the external appearance of the adult *Da* mutant is intriguing, in that it exhibits a teardrop body shape (typical for fish swimming in the middle layer like tuna), instead of a dorsally flattened one (typical for surface swimming fish like medaka). Since our study shows that the activity of *wnt11r* could influence the body shape by regulating cell proliferation and the behavior of muscle progenitor cells, Wnt11r could be one of the crucial factors in evolution and diversity of body shape in fish. Furthermore, the expression of *zic1* (Ohtsuka *et al*., 2004; Sun Rhodes and Merzdorf, 2006; Houtmeyers *et al*., 2013) and *wnt11r* (Marcelle, Stark and Bronner-fraser, 1997; Olivera-Martinez, Thelu and Dhouailly, 2004; Garriock *et al*., 2005; Matsui *et al*., 2005) are strongly conserved among vertebrates, and we hypothesize that Wnt11r-mediated morphogenesis of the somites represents an evolutionarily conserved mechanism that acts across vertebrates.

## Materials and Methods

### Key resource table

#### Animals and transgenic lines

Fish were raised and maintained under standard conditions. All experimental procedures and animal care were performed according to the animal ethics committee of the University of Tokyo. Sex was randomly assigned to experimental groups. Medaka d-rR stain was used as wild-type, the *Da* mutant used in this study was previously described (Ohtsuka *et al*., 2004). The pre-existing transgenic line Tg(*zic1:GFP/zic4:DsRed*) (Kawanishi *et al*., 2013) was used, and the transgenic line Tg(*zic1:GFP/zic4:DsRed*);*Da* was created by crossing *Da* mutants with Tg(*zic1:GFP/zic4:DsRed*). The transgenic line Tg(*zic1:zic1-Myc/zic4:DsRed*);*Da* was generated by modifying the BAC used to generate Tg(*zic1:GFP/zic4:DsRed*) (Kawanishi *et al*., 2013), by replacing the ORF of *GFP* with the ORF of *zic1* containing a sequence for a Myc-tag fused to its C-terminus. To establish the transgenic line, the BAC(*zic1:zic1-Myc/zic4:DsRed)* was co-injected with I-SceI Meganuclease (NEB) into 1-cell stage *Da* embryos, as previously described (Thermes *et al*., 2002).

#### Visualization of actin skeleton of protrusions

The AC-TagGFP2 sequence from the Actin-Chromobody plasmid (TagGFP2) (Chromotek) was cloned into the pMTB vector for mRNA generation. To investigate the actin skeleton of protrusions, cells of Wt and *Da* embryos were mosaically labelled using *Actin-Chromobody-GFP* (*AC-GFP*) mRNA. Embryos were injected at 1-cell stage with 152 ng/µl *membrane-mCherry* mRNA. At 4-cell stage, one cell was injected with 184 ng/µl *AC-GFP* mRNA.

#### Lineage tracing of mesenchymal DM cells

Mosaic labelling of cells was achieved by co-injecting 20 ng/µl pMTB-memb-mTagBFP2 plasmid with Tol2 into a 1-cell stage embryo of the transgenic line Tg(*zic1:GFP/zic4:DsRed*). Embryos were raised until 5 dpf and labelled cells were continuously observed until 8 dpf.

#### Morpholino injection

Microinjections into medaka embryos were performed using 12.5 µM *wnt11r* Morpholino antisense oligonucleotides (MO). Injections were performed into 1-cell stage embryos.

#### Photo-Morpholino mutagenesis

*Wnt11r* Sense-Photo-Morpholino (PhotoMO) and *wnt11r* antisense Morpholino were annealed in a ratio 2:1. Microinjection of 25 µM of the annealed oligonucleotides was performed into 1-cell stage embryos and embryos were raised until 4 ss in the dark. Photo-cleavage was performed using the 10x objective and the DAPI filter of a Keyence BZ-9000 Biorevo microscope (Keyence). Embryos were mounted, dorsal side facing up, in 1 % Methylcellulose in a glass bottom dish (Wako) and illuminated for 30 min. After Photo-cleavage, embryos were dechorionated and raised until the desired stage for subsequent analysis.

#### Injection of human recombinant Wnt11 protein into *Da* mutant somite

To immobilize embryos, embryos from the Tg(*zic1:GFP/zic4:DsRed*);*Da* transgenic line were injected with 25 ng/µl *α-bungarotoxin* mRNA at 1-cell stage. At 18 ss embryos were mounted in 1 % low melting agarose in 1x Yamamoto’s Ringer Solution and oriented with the dorsal side facing upwards. Embryos were injected on top of the 10^th^ somite with a mix containing Dextran Rhodamine (Thermo Fisher) and 1.7 ng hrWnt11 protein (R&D Systems) or BSA (Sigma-Aldrich) and raised to 24 ss, followed by *in vivo* imaging and analysis.

#### KN-93 treatment

Dechorionated 4 ss embryos were treated with 30 μM KN-93 (Wako, Japan) or DMSO (Sigma Aldrich, Germany) in 1x Yamamoto’s Ringer Solution at 28 °C, in the dark until 22 ss was reached.

#### *In vivo* imaging and *in vivo* time-lapse imaging

To immobilize embryos for *in vivo* imaging, embryos were injected at 1-cell stage with 25 ng/µl *α-bungarotoxin* mRNA (Swinburne *et al*., 2015; Lischik, Adelmann and Wittbrodt, 2019). Embryos were mounted in 1 % Low melting agarose (Sigma Adrich) in 1x Yamamoto’s Ringer Solution in a glass-bottomed petri dish (IWAKI) and oriented with the dorsal side facing down. Imaging was performed using a Zeiss LSM 710 confocal microscope system (Zeiss) equipped with an inverted stand and a Zeiss AXIO Observer Z1 and a T-PMT detector. The embryos were positioned with the 5^th^ or 10^th^ somite in the center and images were acquired using a 40x water objective. For the *in vivo* time-lapse imaging, the 10^th^ or 15^th^ somite was positioned in the center, z-stacks were imaged in a 600-sec interval for 10-15 h. Image analysis was performed in Fiji using the “Image Stabilizer” Plugin, the FFT Bandpass filter and the “Draw_arrows” Plugin to draw customized arrows (Li, 2008; Daetwyler, Modes and Fiolka, 2020).

#### Whole mount *in situ* hybridization

Whole mount *in situ* hybridization was performed as previously described with the following modifications (Takashima *et al*., 2007). Embryos were fixed in 4 % PFA/1.5x PTW at 4 °C, overnight. Hybridization was performed at 65 °C, overnight. Samples were treated with alkaline-phosphatide anti-DIG-AP Fab fragments (1:2000, Roche). Signals were developed using 4-nitro blue tetrazolium chloride (NBT, Roche) and 5-bromo-4-chloro-3-indolyl phosphate (BCIP, Roche).

#### Whole-mount immunohistochemistry

Embryos were fixed in 4 % PFA/PBS for 2 h at room temperature or at 4 °C overnight. Samples were permeabilized with 0.5 % TritonX-100 (Wako) in 1x PBS for 1-2 h and blocked in blocking solution (2 % BSA (Sigma-Aldrich), 1 % DMSO, 0.2 % TritonX-100 in 1x PBS) for 2-4 h at room temperature. Samples were incubated with respective primary antibody diluted in blocking solution at 4 °C, overnight. After an additional 4 h blocking step, samples were incubated with respective secondary antibodies diluted in blocking solution, at 4 °C, overnight. Samples were stored in 1x PBS at 4 °C until imaging.

#### Antibodies

| Primary antibody | Species | Concentration | Company |
| --- | --- | --- | --- |
| anti-Pax3/7 (DP312) | mouse | 1:100 | Davis et al., 2001 |
| anti-Ph3 | rabbit | 1:200 | Millipore, 06-570 |
| anti-GFP | rabbit | 1:200 | Clontech, 632593 |

| Secondary antibody | Species | Concentration | Company |
| --- | --- | --- | --- |
| anti-mouse Alexa 555 | goat | 1:500 | Thermo Fisher, A-21422 |
| anti-rabbit Alexa 488 | donkey | 1:500 | Life Technnologies, A-11008 |

#### Vibratome sectioning

Samples were mounted in 4 % agarose in 1x PBS. 40 μm or 200 µm sections were obtained by a Vibratome (Leica, Vibratome). The sections were mounted on a glass slide (Matsunami) in 60 % glycerol (Merck, Wako) and stored at 4 °C until imaging. Images were acquired using the 40x water objective of a Zeiss LSM 710 confocal microscope.

#### Histological sections

Dechorionated embryos were fixed in Bouin’s solution overnight, followed by a gradual dehydration using ethanol. Samples were embedded in Technovit 7100 (Heraeus Kulzer) and sectioned into 5-6 µm thick sections. Sections were stained with hematoxylin (Wako) and imaged using the 1.6x objective of a Leica M165 FC fluorescent stereo microscope.

#### Image processing and statistical analysis

Image processing was performed with the image processing software Fiji. The 3D-recreation of *in vivo* imaging date was created using FluoRender (Wan *et al*., 2009). Measurements of morphological features (distance between myotome tips, area of cross-section of dorsal somites, diameter of myofibers, distance between dorsal somite tip and the tip of neural tube, somite height) was performed by averaging the analysis of the feature from 3 consecutive Y planes. RStudio was used for the statistical analysis and representation of the data. In bar plots mean and error limits, defined by the standard deviation, are indicated. In box plots median first and third quantiles are indicated. Statistical significance was determined by unpaired t-tests, a p-value p < 0.05 was considered as significant. In the figure legends sample size (n) and number of individuals used in the experiment are stated. Sample sizes were not predetermined using statistical methods, but the sample sizes used are similar to those generally used in the field. To compare experimental groups, the allocation was performed randomly, without blinding.

#### Isolation of dorsal and ventral somite cells for ATAC-seq and RNA-seq

Yolk and head region were removed from 22 ss Tg(*zic1:GFP/zic4:DsRed*) embryos and embryos were incubated with 10 mg/ml pancreatin (Wako) at room temperature for 5-10 min. Epidermis and intestinal tissues were removed and somites were isolated from the neural tube. Cells were dissociated in 0.5% (w/v) Trypsin (Nacalai Tesque) at 37 °C for 10 min, the dissociation was stopped by adding the same volume of 15% (v/v) FBS / Leiboviz’s L-15 (Life Technologies). Dissociated cells were washed with PBS, and sorted into GFP positive (dorsal) and negative (ventral) cells using FACSAria III (BD Biosciences). Dead cells were detected by Propidium iodide (Life Technologies) and removed.

#### RNA-seq

Total RNA was extracted from sorted somite cells using RNeasy Mini kit (Qiagen). mRNA was enriched by poly-A capture and mRNA-seq libraries were generated using KAPA Stranded mRNA-seq Kit (KAPA Biosystems). Libraries were generated from two biological replicates, and sequenced using the Illumina HiSeq 1500 platform.

#### RNA-seq data processing

The sequenced reads were pre-processed to remove low-quality bases and adapter derived sequences using Trimmomatic v0.32 (Bolger, Lohse and Usadel, 2014), and aligned to the medaka reference genome (HdrR, ASM223467v1) by STAR (Dobin *et al*., 2013). Reads with mapping quality (MAPQ) larger than or equal to 20 were used for the further analyses.

#### ATAC-seq

ATAC-seq was performed as previously described (Buenrostro *et al*., 2013) with some modifications. Approximately 4,000 sorted somite cells were used for each experiment. After washing with PBS, cells were resuspended in 500 μl cold lysis buffer (10 mM Tris-HCl pH 7.4, 10 mM NaCl, 3 mM MgCl_2_, 0.1% Igepal CA-630), centrifuged for 10 min at 500 g, supernatant was removed. Tagmentation reaction was performed as described previously (Buenrostro *et al*., 2013) with Nextera Sample Preparation Kit (Illumina). After tagmented DNA was purified using MinElute kit (Qiagen), two sequential PCRs were performed to enrich small DNA fragments. First, a 9-cycle PCR was performed using indexed primers from Nextera Index Kit (Illumina) and KAPA HiFi HotStart ReadyMix (KAPA Biosystems), amplified DNA was size selected for a size less than 500 bp using AMPure XP beads (Beckman Coulter). A second 7-cycle PCR was performed using the same primers as for the first PCR. PCR product was purified by AMPure XP beads. Libraries were generated from two biological replicates, and sequenced using the Illumina HiSeq 1500 platform.

#### ChIPmentation

Yolk and head region were removed from 22 ss Tg(*zic1:zic1-Myc/zic4:DsRed*);*Da* embryos, followed by an incubation with 10 mg/ml pancreatin (Wako) at room temperature for 5-10 min. Epidermis and intestinal tissues were removed and somites were isolated from the neural tube. ChIP was performed as previously described with the following modifications (Nakamura *et al*., 2014). Isolated somites were fixed with 1% formaldehyde for 8 min at room temperature then quenched by adding glycine (200 mM final) and incubating on ice for 5 min. After washing with PBS, cell pellets were stored at −80 ℃. Approximately 1.8×10^6^ cells were thawed on ice, suspended in lysis buffer (50 mM Tris-HCl (pH 8.0), 10 mM EDTA, 1% SDS, 20 mM Na-butyrate, complete protease inhibitors, 1 mM PMSF) and sonicated 10 times using a Sonifier (Branson) at power 5. Chromatin lysates were collected by centrifugation and diluted 10-fold with RIPA ChIP buffer (10 mM Tris-HCl (pH 8.0), 140 mM NaCl,1 mM EDTA, 0.5 mM EGTA, 1% Triton X-100, 0.1% SDS, 0.2% sodium deoxycholate, 20 mM Na-butyrate, complete protease inhibitors, 1 mM PMSF) followed by an incubation with antibody/protein A Dynabeads (Invitrogen) complex at 4 °C, overnight, while rotating. Immunoprecipitated samples were washed three times with RIPA buffer (10 mM Tris-HCl (pH 8.0), 140 mM NaCl,1 mM EDTA, 0.5 mM EGTA, 1% Triton X-100, 0.1% SDS, 0.2% sodium deoxycholate) and once with TE buffer. After the washing steps 150 μl of Tris-HCl was added.

Library preparation for ChIPmentation was performed as previously described (Schmidl *et al*., 2015) with the following modifications. 24 μl of Tagmentation reaction mix (10 mM Tris-HCl pH8.0, 5 mM MgCl_2_, 10%(v/v) N,N-dimethyl formamide) and 1 μl of Tagment DNA Enzyme from Nextera Sample Preparation Kit (Illumina) were added to the DNA-beads complex and incubated for 70 sec at 37 °C. 150 μl ice-cold RIPA buffer was added and incubated for 5 min on ice. The DNA-beads complex was washed with RIPA buffer and TE buffer, suspended in 50 μl lysis buffer and 3 μl of 5 M NaCl, and incubated at 65 °C, overnight. The sample was incubated for 2 h with 2 μl of 20 mg/ml ProteinasK (Roche), and subjected to AMPure XP beads (Beckman Coulter) purification. The library was amplified by 18-cycle PCR using indexed primers from Nextera Index Kit (Illumina) and KAPA HiFi HotStart ReadyMix (KAPA Biosystems). For the input chromatin, tagmentation reaction was performed after DNA purification. Libraries were generated from two biological replicates, and sequenced using the Illumina HiSeq 1500 platform.

#### ChIPmentation and ATAC-seq data processing

The sequenced reads were pre-processed to remove low-quality bases and adapter derived sequences using Trimmomatic v0.32 (Bolger, Lohse and Usadel, 2014) and aligned to the medaka reference genome (HdrR, ASM223467v1) by BWA (Li and Durbin, 2009). Reads with mapping quality (MAPQ) larger than or equal to 20 were used for the further analyses. MACS2 (version 2.1.1.20160309) (Zhang *et al*., 2008) was used to call peaks and generate signals per million reads tracks using following options; ChIPmentation: -g 600000000 -B -- SPMR --keep-dup 2, ATAC-seq: --nomodel --extsize 200 --shift -100 -g 600000000 -q 0.01 - B --SPMR.

For ChIPmentation, peak regions called by two biological replicates were used as reliable peaks.

#### Motif analyses of Zic1 ChIPmentation peaks

Motifs enriched at reliable ChIPmentation peaks were analyzed by findMotifsGenome command of HOMER (Heinz *et al*., 2010) using default parameters.

#### Identification of Zic1 target genes

Differentially expressed genes were identified using DESeq2 (padj < 0.01). Each reliable ChIP peak was associated to the nearest TSS, and the gene was defined as Zic1-target gene if the distance between the peak and the TSS was closer than 50 kb.

#### Gene ontology and pathway analyses

The gene ontology enrichment analyses and pathway enrichment analyses were performed using the Gene Ontology Resource (Ashburner *et al*., 2000; Gene Ontology Consortium, 2021).

#### RT-PCR of cDNA generated from embryonic tails

To investigate the gene expression in tails of embryos, tails were dissected anterior from the first somite. Ten tails were pooled together and RNA was isolated using Isogen (Nippon Gene). RNA was purified using the RNeasy Mini kit (Qiagen) and reverse transcribed to cDNA using the Super Script III Kit (Invitrogen). RT-PCR was performed using the Thunderbird Sybr qPCR Mix (Toyobo) following manufacturer’s instructions and run in the Agilent Mx3000P qPCR System (Agilent). Normalization of relative quantities was performed against *gapdh* expression, followed by analysis with excel and RStudio.

## Acknowledgments

We thank the members of the Takeda laboratory for constructive feedback and discussions on the project. We are greatful for Y. Yamagichi and M. Funato for fish husbandry. This work was supported by Japan Society for the Promotion of Science (JSPS) KAKENHI Grant Numbers JP15H05859 (H.T.), JP19K23741 (T.K.) and JP18K14620 (R.N.) and Japan Science and Technology Agency CREST Grant Number JPMJCR13W3 (H.T.).

---

**Figure 1 - figure supplement 1:**
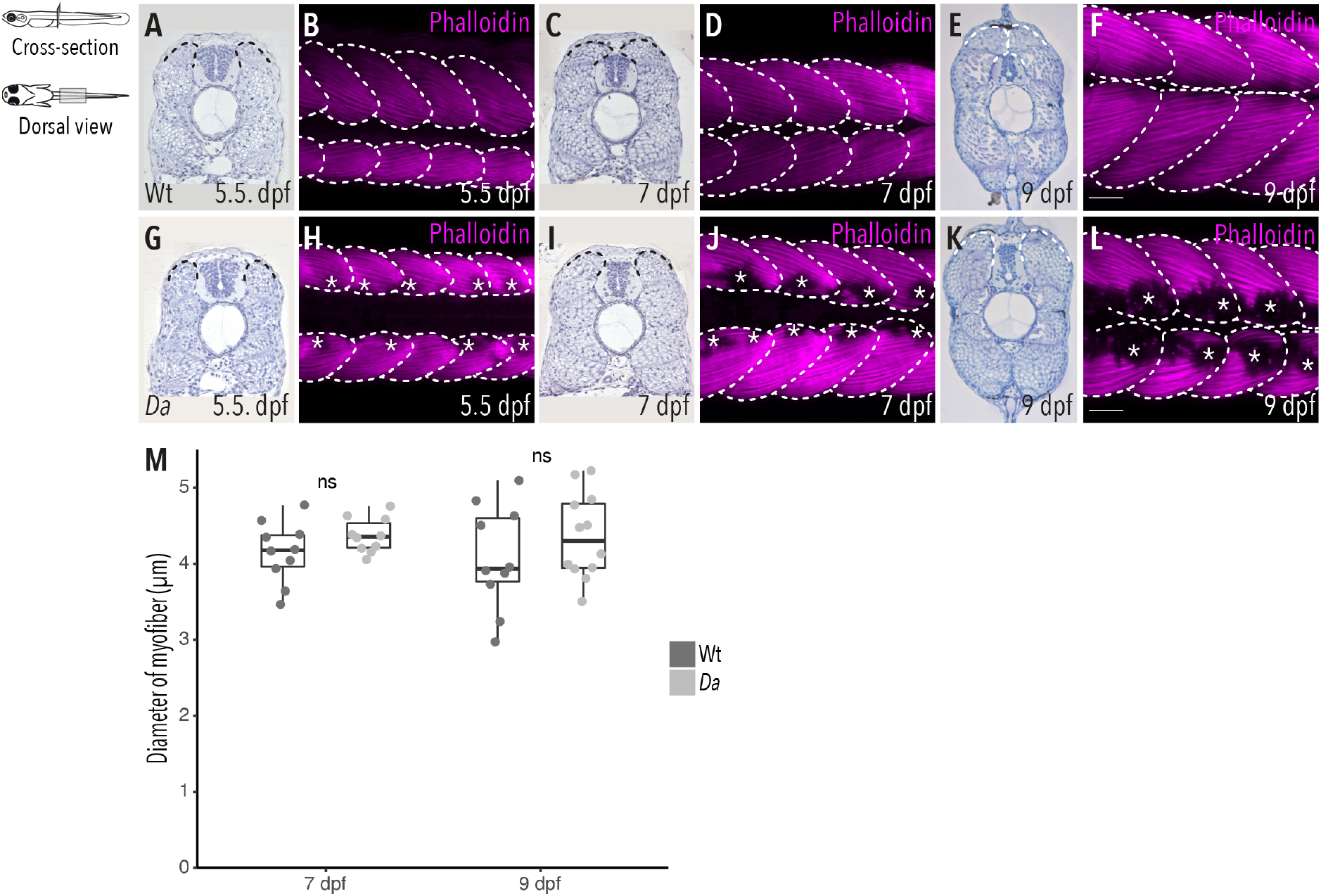
The ventralized epaxial myotome of the *Da* mutant fails to extend sufficiently to cover the neural tube at the end of embryonic development. (A, C, E,) Hematoxylin staining of cross-sections from Wt and (G, I, K) *Da* mutant embryos and dorsal view of whole-mount Phalloidin (magenta) immunostaining to label the myotome of Wt (B, D, F) and *Da* mutant embryos (H, J, L). (M) Analysis of diameter of myofibers in dorsal myotome of 7 dpf embryos (n = 10 myofibers from 10 somites of 5 Wt embryos, n = 10 myofibers from 10 somites of 5 *Da* embryos) and 9 dpf embryos (n = 10 myofibers from 10 somites of 5 Wt embryos, n = 12 myofibers from 12 somites of 6 *Da* embryos) (median, first and third quartiles, ^ns^P_7 dpf_ = 0.15, ^ns^P_9 dpf_ = 0.31). Anterior = left, scale bar = 50 μm.

**Figure 2 - figure supplement 1:**
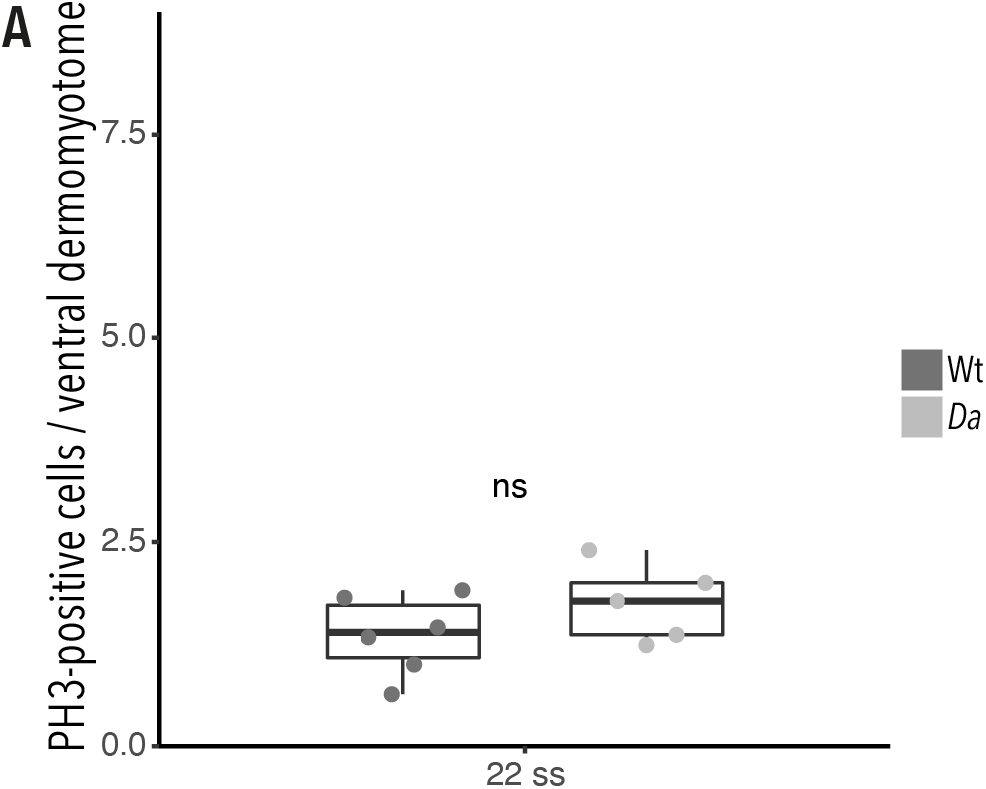
Difference in proliferative activity is not observed between Wt and *Da* ventral DM. (A) Quantification of PH3-positive cells in the ventral DM of 22 ss embryos (n = 66 somites from 6 Wt embryos, n = 51 somites from 5 *Da* embryos) (median, first and third quartiles, ^ns^P = 0.2).

**Figure 3 - figure supplement 1:**
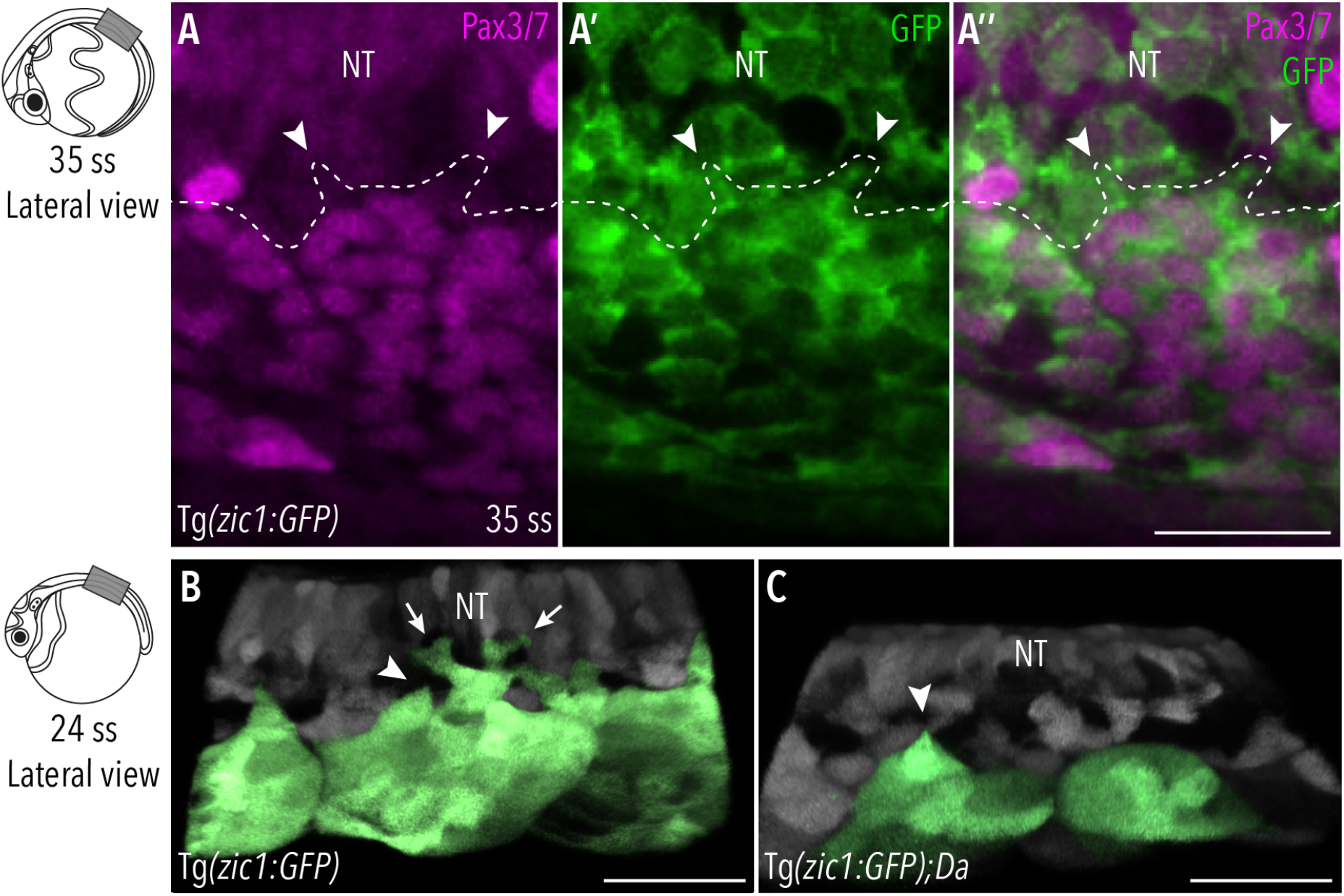
Protrusion forming cells at the tips of the dorsal somites are DM cells. (A-A’’) Dorsal view of immunostaining performed on Tg(*zic1:GFP*) embryos at 35 ss. GPF labels Zic1-positive dorsal somite cells (green), Pax3/7 labels DM cells (magenta). Arrowheads indicate protrusions, scale bar = 50 μm. (B-C) Lateral view of 3D-reconstractions of dorsal somites (green) from (B) Tg(*zic1:GFP*) and (C) Tg(*zic1:GFP*);*Da* embryos at 24 ss. 10^th^ somite is positioned in center, small and large protrusions are exemplary labeled by arrowheads and arrows respectively. Scale bar = 25 μm. Anterior = left, dorsal = up, NT = neural tube.

**Figure 3 - figure supplement 2:**
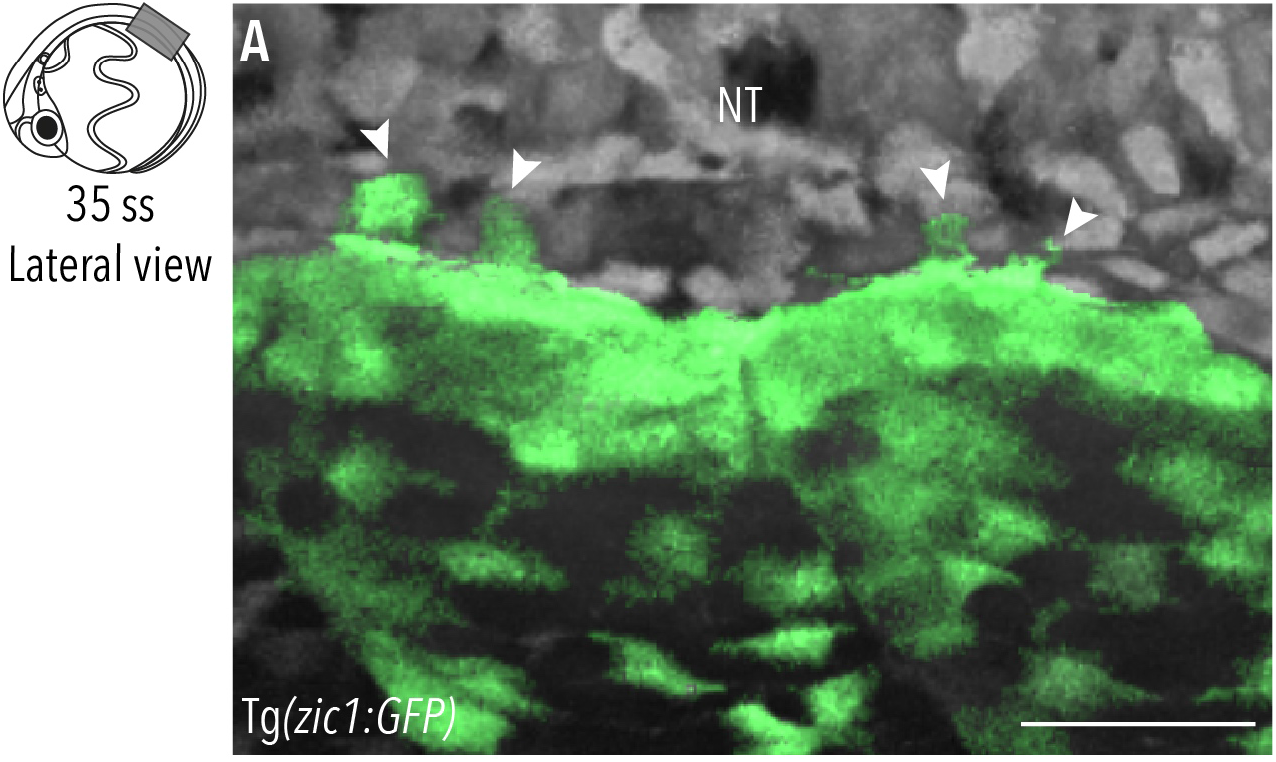
Dorsal DM cells form protrusions throughout dorsal somite extension. (A) Lateral view of 3D-reconstractions of dorsal somites (green) from Tg(*zic1:GFP*) at 35 ss 5^th^ somite is positioned in center, large protrusions are indicated with arrowheads. Anterior = left, dorsal = up, NT = neural tube, scale bar = 50 μm

**Figure 4 - figure supplement 1:**
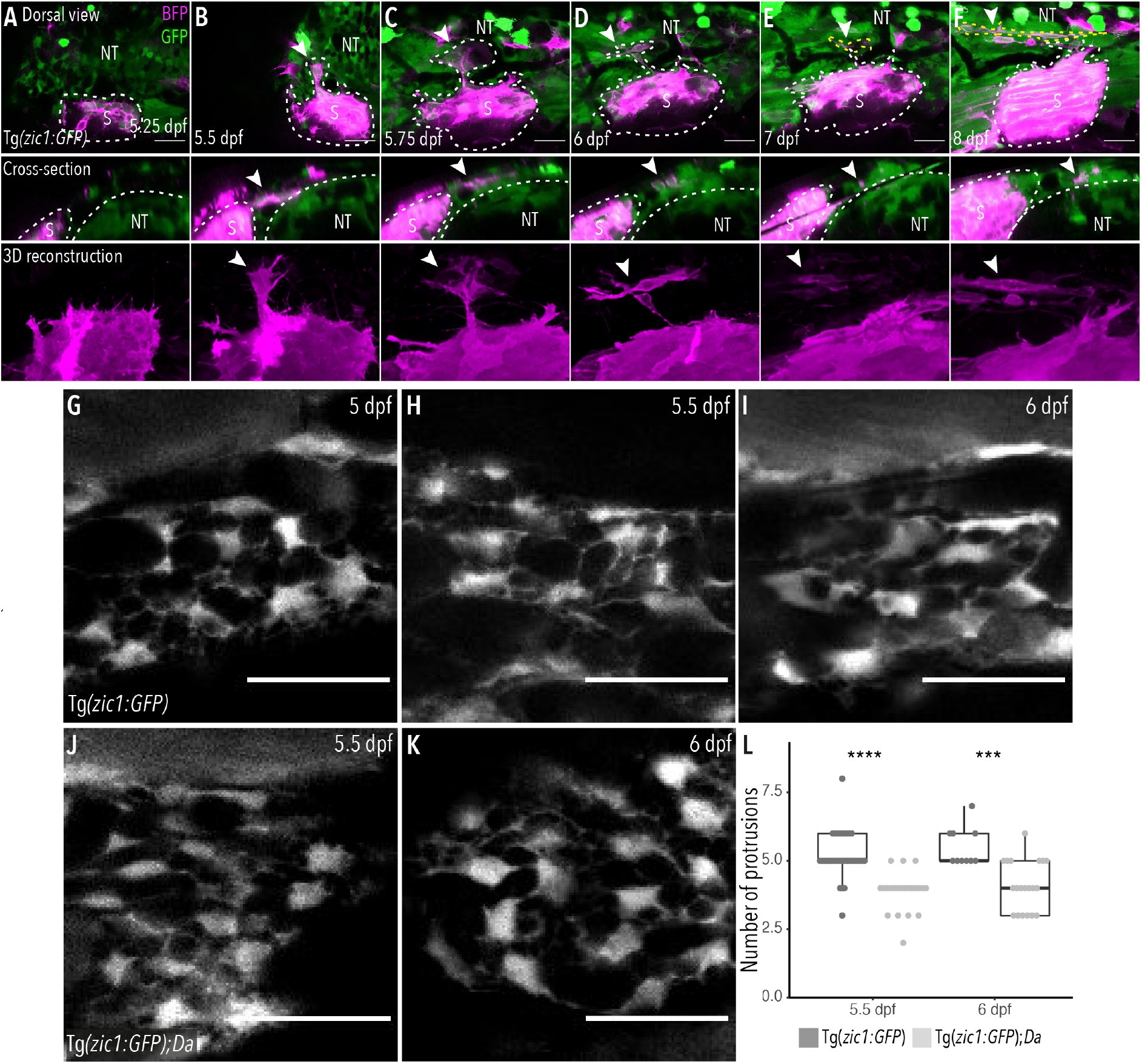
Mesenchymal DM cells originate from dorsal somites. (A-F) Lineage tracing experiment to clarify the origin of mesenchymal DM cells. Dorsal somite cells express BFP (magenta). (B-F) Arrowheads indicate DM cell originating from labelled somite which (B-C) extends dorsally and (D) delaminates, becoming a mesenchymal DM cell. Somites are outlined in white, mesenchymal DM cell is outlined in yellow (E, F). (G-K) Maximum projections of mesenchymal DM cells of Tg(*zic1:GFP*) at (G) 5 dpf, (H) 5.5 dpf and (I) 6 dpf and Tg(*zic1:GFP*);*Da* at (J) 5.5 dpf and (K) 6 dpf. (L) Quantification of protrusions of mesenchymal DM cells at 5.5 dpf (n = 40 cells from Tg(*zic1:GFP*), n = 22 cells from Tg(*zic1:GFP*);*Da*) and 6 dpf (n = 10 cells from Tg(*zic1:GFP*), n = 20 cells from Tg(*zic1:GFP*);*Da*)(median, first and third quartiles, ****P_5.5 dpf_ = 4.4e-07, ***P_6 dpf_ = 0.00012,). Anterior = left, NT = neural tube, scale bar = 25μm.

**Figure 4 - figure supplement 2:**
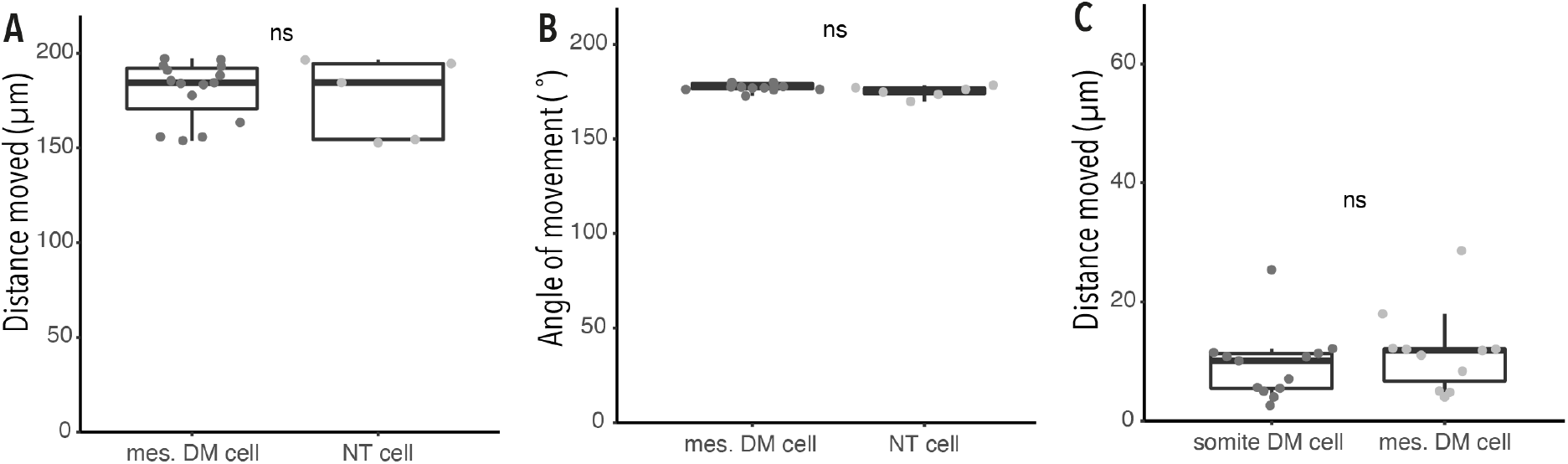
Mesenchymal DM cells do not migrate significantly. (A) Analysis of distance moved relative to imaging field by mesenchymal DM cells (mes. DM cell, n = 15 DM cells) and cells of the neural tube (NT cell, n = 5 NT cells) during 13.3 h *in vivo* imaging of 4.5 dpf Tg(*zic1:GFP*) embryo (video 3) (median, first and third quartiles, ^ns^P = 0.73). (B) Analysis of angle moved relative to imaging field by mesenchymal DM cells (n = 15 DM cells) and cells from the neural tube (n = 5 NT cells) during 13.3 h *in vivo* imaging of 4.5 dpf Tg(*zic1:GFP*) embryo (video 3) (median, first and third quartiles, ^ns^P = 0.1). (C) Analysis of distance moved relative to imaging field by DM cells from the tip of the dorsal somite (somite DM cell, n = 11 somite DM cells) and mesenchymal DM cells (mes. DM cells, n = 13 mesenchymal DM cells) during 15 h *in vivo* imaging of 5.5 dpf Tg(*zic1:GFP*) embryo (video 4) (median, first and third quartiles, ^ns^P = 0.4).

**Figure 5 - figure supplement 1:**
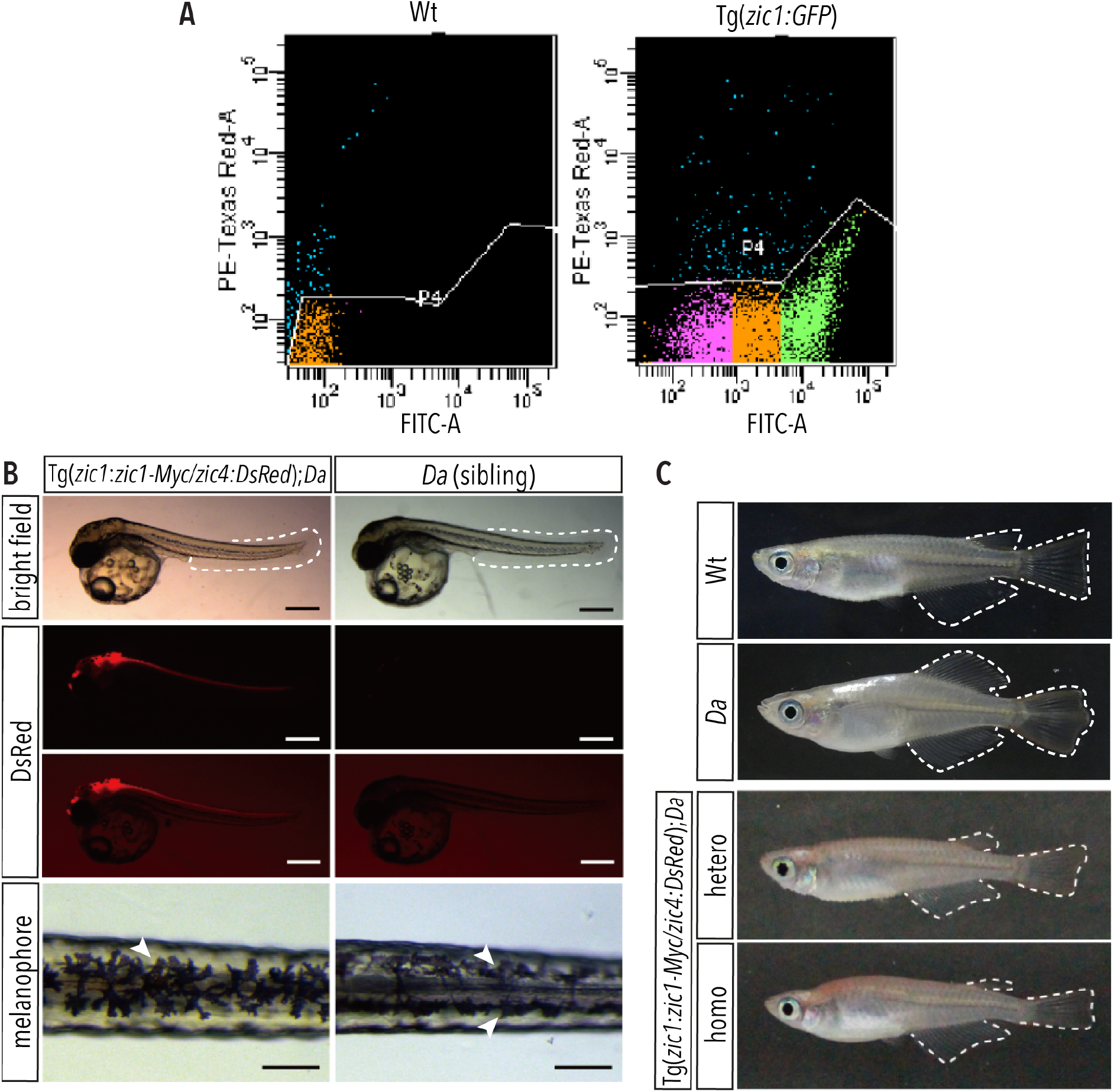
In Tg(*zic1:zic1-Myc*);*Da* fish, the ventralized trunk phenotype is rescued. (A) A representative flow cytometry plot of GFP expression in somite cells. Cells in green and magenta regions were collected as GFP positive and negative cells, respectively. PE-Texas Red-A indicates dead cells by PI staining. FITC shows GFP signal intensity. (B-C) *Da* phenotype is rescued in the transgenic line Tg(*zic1:zic1-Myc*);*Da*. DsRed is expressed in the neural tube and dorsal somites, mimicking the endogenous expression of *zic1* and *zic4* (scale bar = 500 μm). *Da* mutant embryos display two rows of melanophores on the dorsal midline of their trunk (arrowheads, scale bar = 100 μm). In Tg(*zic1:zic1-Myc*);*Da* embryos, as in Wt embryos, a single row of melanophores (arrowhead) on the dorsal midline is found. Ventralized phenotypes of adult *Da* mutants (including body shape, pigmentation, fin shape (outlined)) are rescued in Tg(*zic1:zic1-Myc*);*Da* adult fish. (C) Images of Wt and *Da* adult medaka were first shown in Figure 1B,D.

**Figure 5 - figure supplement 2:**
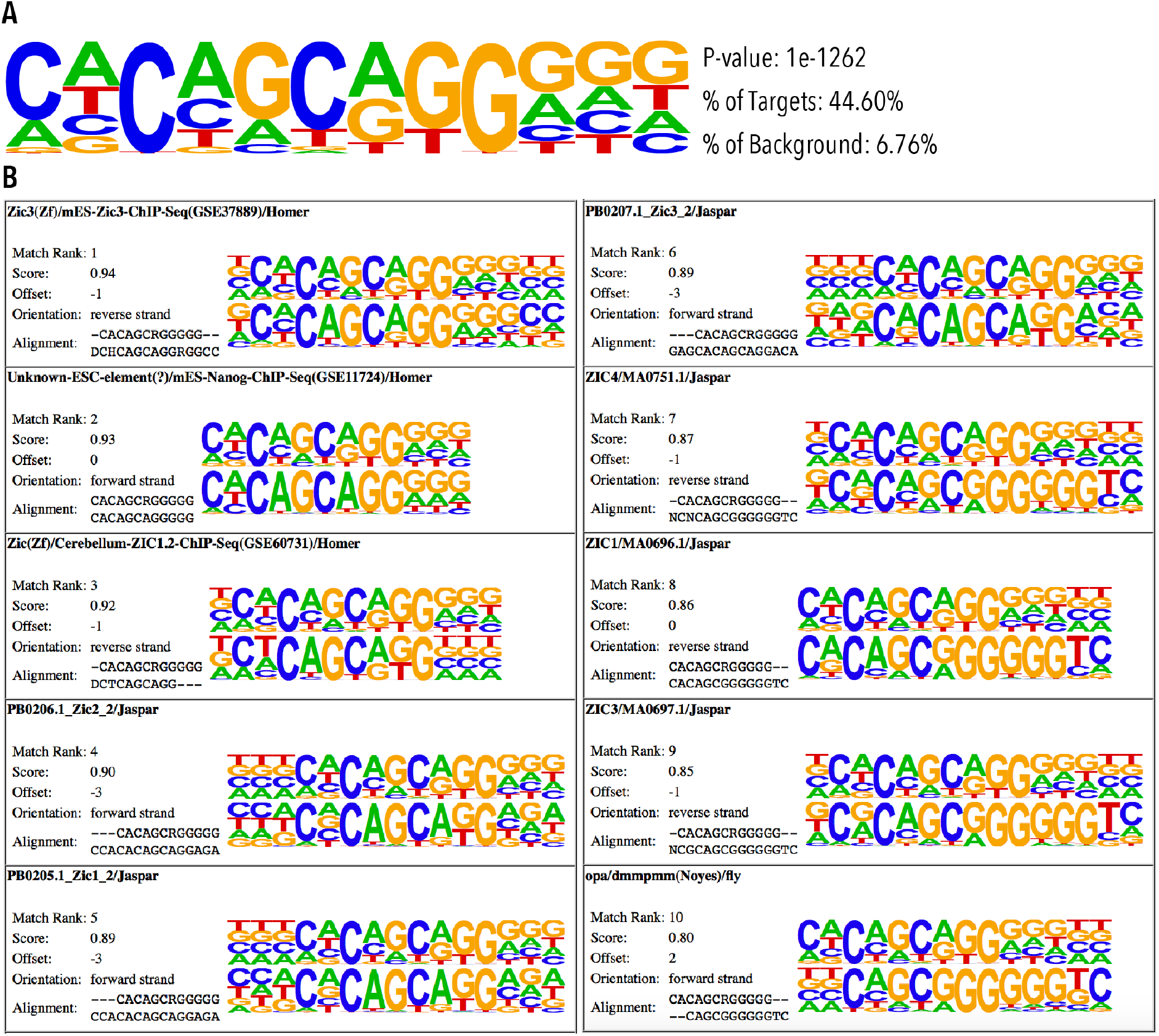
Common ZIC family DNA motifs are enriched in Zic1 peaks. (A) Top DNA motif enriched in Zic-Myc ChIPmentation peaks identified by HOMER. (B) Known motifs matched to DNA motif in (A). For each match, the upper motif, identified in this study (A), was compared to a known motif (lower motif). Matches were analyzed by HOMER.

**Figure 6 - figure supplement 1:**
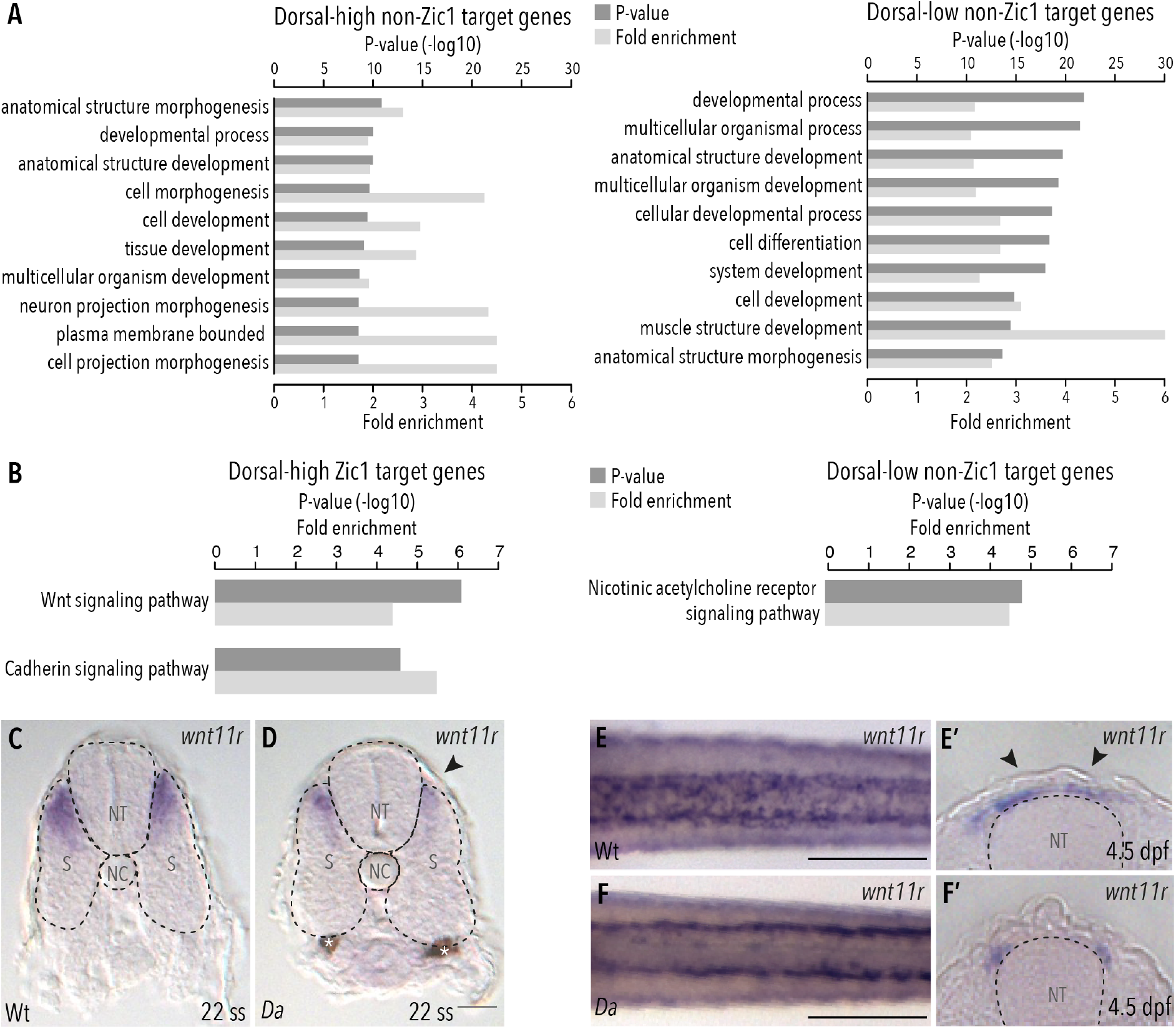
Wnt signaling pathway is enriched in dorsal-high Zic1 target genes. (A) GO analysis of dorsal-high and dorsal-low non-Zic1 target genes. (B) Pathway enrichment analysis of dorsal-high Zic1 target genes and dorsal-low non-Zic1 target genes. (C-D) Sections of the trunk of whole-mount *in situ* hybridization against *wnt11r* performed on 22 ss (C) Wt and (D) *Da* mutant embryos. The expression of *wnt11r* is reduced in the dorsal somites of the *Da* mutant (arrowhead). Asterisks mark melanophores, scale bar = 38 μm. (E-F) Whole-mount *in situ* hybridization against *wnt11r* on (E-E’) Wt and (F-F’) *Da* embryos at 4.5 dpf (dorsal view: E,F, cross-section: E’, F’). In Wt *wnt11r* expression is further restricted to the dorsal most part of the somites (E’) and the mesenchymal DM cells (E-E’, arrowheads). The expression of *wnt11r* is reduced in the dorsal somites of the *Da* mutant (F-F’). Scale bar = 100 μm Dorsal = up, NC = notochord, NT = neural tube, S = somite, scale bar = 38 μm, sections = 40 μm.

**Figure 7 - figure supplement 1:**
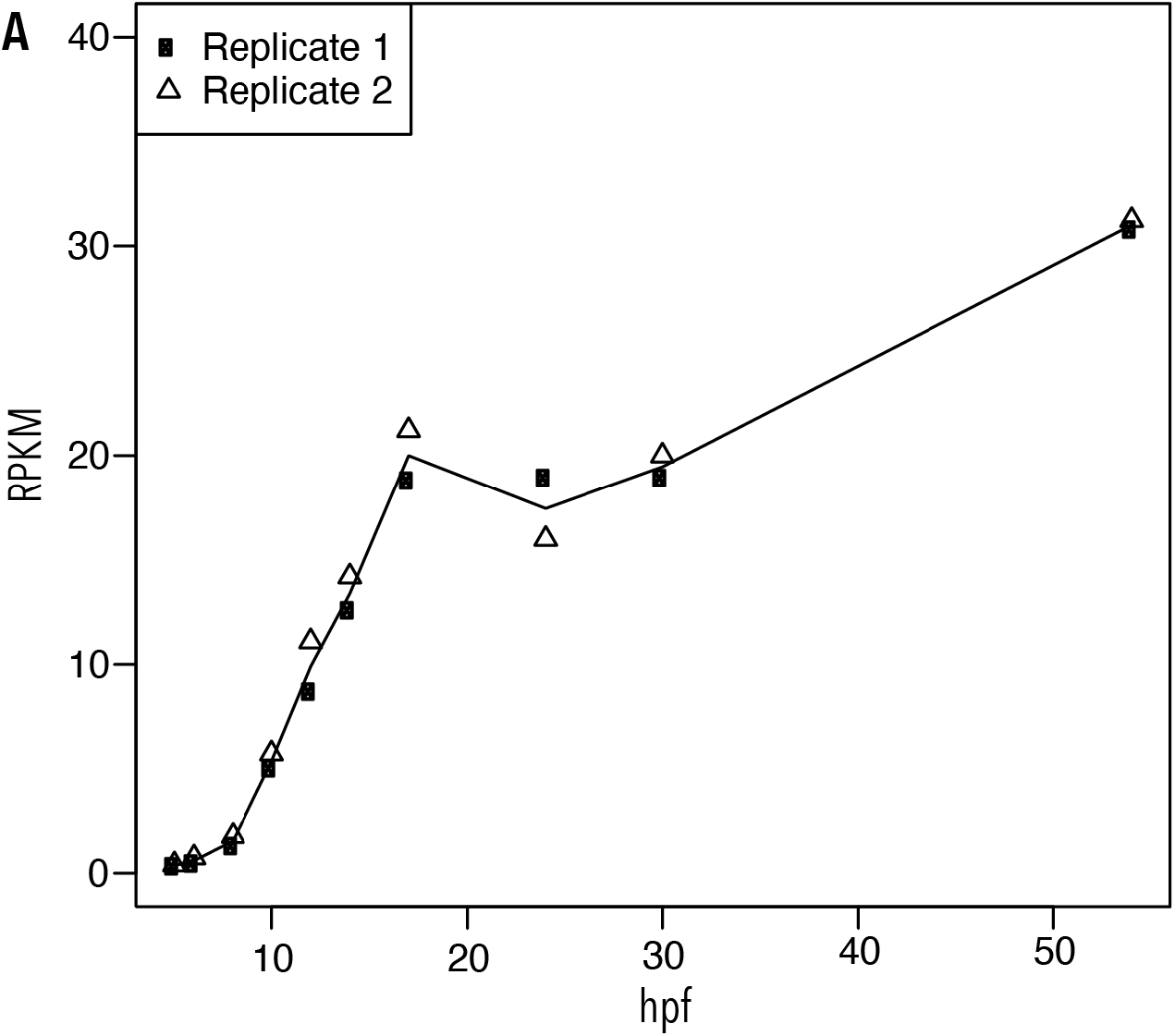
*Wnt11r* expression starts before gastrulation and increases with proceeding development. (A) Reads per kilo base per million mapped reads (RPKM) of *wnt11r* at 5 hpf (hours post fertilization, stage 9), 6 hpf (stage 10), 8 hpf (stage 11), 10 hpf (stage 12), 12 hpf (stage 13, onset of gastrulation), 14 hpf (stage 14), 17 hpf (stage 15, gastrulation competed), 24 hpf (stage 18), 30 hpf (6 ss, stage 21) and 54 hpf (24 ss, stage 27).

Supplementary table 1: Gene expression profiles and distances to nearest Zic1 ChIP peak

Supplementary table 2: Full list of GO terms enriched in dorsal-high Zic1 target genes

Supplementary table 3: Full list of GO terms enriched in dorsal-low Zic1 target genes

